# The toll-like receptor signalling pathway is altered in iPSC-derived cortical networks from people with bipolar disorder

**DOI:** 10.64898/2026.06.09.731031

**Authors:** Bruna Panizzutti, Chiara Cristina Bortolasci, Megan Ellis, Briana Spolding, Courtney Swinton, Trang T. T. Truong, Zoe Shih-Jung Liu, Damián Hernández, Greg Roebuck, Ajeet B. Singh, Bruno Agustini, Robson Zazula, Ana C. Andreazza, Dana El Soufi El Sabbagh, Hyunjin Jeong, Olivia M. Dean, Jee Hyun Kim, Michael Berk, Ken Walder

**Affiliations:** Deakin University, School of Medicine, IMPACT, The Institute for Mental and Physical Health and Clinical Translation, Geelong, Australia; Florey Institute of Neuroscience and Mental Health, Parkville, Australia; Barwon Health, Geelong, Australia; Federal University for Latin America Integration, School of Medicine, Foz do Iguacu, Parana, Brazil; Department of Pharmacology & Toxicology and Psychiatry, Mitochondrial Innovation Initiative (MITO2i), University of Toronto, Toronto, ON, Canada

**Author notes:** Corresponding author: Bruna Panizzutti, BSc, PhD Research Fellow The Institute for Mental and Physical Health and Clinical Translation (IMPACT) School of Medicine Deakin University, Geelong, Victoria, Australia, 3216. Corresponding author: Prof. Ken Walder.

**Keywords:** Bipolar Disorder, mania, depression, psychiatry, neuroscience, iPSC-derived brain cells, Toll-like signalling pathway, TLR4

## Abstract

**Background:** Induced pluripotent stem cell (iPSC)-derived brain cells are widely utilized as *in vitro* models for several neuropsychiatric disorders, as they retain the donor’s genetic profile, offering a unique opportunity to study living human brain cells and perform controlled experimental manipulations. In this study, we conducted whole transcriptome sequencing of cortical networks (co-cultures of neurons and astrocytes) derived from 12 participants with bipolar disorder (BD) and 12 participants without a history of mental health disorders. We aimed to identify new molecular mechanisms underlying the pathophysiology of bipolar disorder.

**Methods:** iPSCs were generated by reprogramming peripheral blood mononuclear cells using episomal vectors. They were then differentiated into neural progenitor cells and matured into cortical networks that express markers of neurons and astrocytes. Whole transcriptome data were obtained using the Illumina NovaSeq X sequencing platform.

**Results:** Differential expression analysis was performed using DESeq2 in R, and the identified genes were used for gene set enrichment analysis, which identified 191 enriched pathways in BD. Of these, the toll-like signalling pathway, which is downregulated in BD, was further investigated.

**Conclusion:** Our results suggest a profound immune dysregulation in BD, particularly highlighting the immune system’s role as a complex signalling network.

## Introduction

Twin and family studies estimate the heritability of bipolar disorder (BD) at 60-80%^1,2^, and genome-wide association studies indicate that common genetic variants account for around 20% of BD phenotypic variance^3^. However, the specific biological mechanisms involved in the pathophysiology of BD remain poorly understood. Additionally, the inaccessibility of living brain tissue and the lack of suitable *in vitro* and *in vivo* models make donor-derived induced pluripotent stem cells (iPSCs) an attractive tool for studying fundamental cellular processes in human brain cells that preserve the genetic background of the donors.

The advent of induced pluripotent stem cell (iPSC) technology has revolutionised neuropsychiatric research by enabling the generation of patient-derived brain cells. These models provide direct access to living human brain cells, allowing the study of early neurodevelopmental processes, cellular excitability, and drug responsiveness in a controlled *in vitro* setting and comparison to control stem cells. iPSC-derived neurons have been vital in identifying alterations in neurodevelopmental trajectories, hyperexcitability, and disrupted neuroplasticity^4–6^, all of which are implicated in BD pathophysiology. Moreover, these models facilitate the investigation of pharmacological interventions, including treatment responsiveness, offering a robust platform for hypothesis generation and precision medicine approaches in BD.

In this study, we aim to investigate the transcriptional changes associated with BD using an *in vitro* model of iPSC-derived neurons and astrocytes in conjunction with whole transcriptome sequencing to uncover novel molecular mechanisms that contribute to the pathophysiology of BD.

## Materials & Methods

### Participant recruitment

Potential participants with BD were identified from a database of participants who agreed to be contacted for research opportunities or via referral from clinicians at The Geelong Clinic and invited to participate. Control participants were recruited from the community via advertisement flyers and, after a telephone screen, invited to participate in the study. Recruitment occurred between September 2019 and December 2021. Trained researchers conducted the study assessments, which included the Structured Clinical Interview for DSM-5^7^ (SCID-5-RV version 1.0.0; patient or non-patient versions) and the BRFSS Adverse Childhood Experience (ACE) module. A phlebotomist performed cubital blood sample collection. The psychiatrist overseeing the care of the participants with BD was then contacted to complete the lithium response phenotype scale – the Alda scale^8^. Scores of 7 or above were deemed lithium responders, while scores below 7 were lithium non-responders. Inclusion criteria included: participants aged 18 or over, who fulfilled the DSM-V diagnostic criteria for BD (cases) or no mental health disorders (controls) and had the capacity to consent to the study and comply with study procedures as ascertained by the researcher. Exclusion criteria included concurrent diagnosis of other mental health disorders rather than BD, known or suspected clinically relevant systemic medical disorder, females who were pregnant or lactating, inability to comply with either the requirements of informed consent or the treatment protocol and current enrolment in a clinical trial. This study received human ethics approval from Barwon Health Hospital (17/205) and Deakin University (2018-316). Clinical and demographic information is in Tables 1.

**Table 1.** Descriptive statistics of participants information. Age and BMI were expressed as mean (standard deviation).

|  | *Bipolar disorder group*<br>*(n=12)* | *No mental health disorders group*<br>*(n=12)* | *p-*<br>*value* |
| --- | --- | --- | --- |
| *Age* | 48.17 (14.84) | 44.83 (11.11) | 0.539 |
| *BMI* | 27.17 (4.15) | 23.91 (5.02) | **0.018** |
| *Females* | 9 (75%) | 11 (91.6%) |  |
|  | Lithium Responders (n=5) | Lithium non -responders (n=6) |  |
| *Age* | 55.80 (14.64) | 41.50 (14.18) | 0.135 |
| *BMI* | 28.20 (4.91) | 25.83 (3.71) | 0.386 |
| *Females* | 3 (60%) | 6 (110%) |  |

### Peripheral blood mononuclear cells (PBMCs) isolation

Blood samples were collected in EDTA vacutainer tubes (Becton Dickinson, NJ, US) from 11 healthy adults (plus ATCC5.1 cell line reprogrammed from commercially available PBMCs) and 12 participants with BD. Blood was diluted 1:2 in wash buffer (PBS/2% FBS/2mM EDTA), layered into SepMate™-15 tubes with Lymphoprep (both, StemCell Technologies, Canada), and centrifuged at 1200g for 10 mins. The PBMC layer was transferred, washed with wash buffer, and centrifuged at 300g for 10 mins. The pellet was rinsed with ice-cold Milli Q water to remove red blood cells, then washed in buffer and centrifuged again at 300g for 10 mins. The cells were cryopreserved in FBS +10%DMSO.

### PBMCs reprogramming to iPSCs

PBMCs were cultured for 7 days in StemSpan™ SFEM II with erythroid expansion supplement (StemCell Technologies, Canada). Reprogramming was performed using the Epi5™ Episomal iPSC Reprogramming Kit and Neon™ Transfection System Kit (Invitrogen, Thermo Fisher Scientific, MA, US). Transduced cells were plated on Geltrex™-coated dishes (Gibco, Thermo Fisher Scientific, MA, US) and maintained in mTeSR Plus medium (StemCell Technologies, Canada). Single iPSC colonies were manually isolated, expanded via enzymatic passaging with ReLESR (StemCell Technologies, Canada), and cultured in Geltrex-coated flasks with mTeSR Plus medium. Stocks were cryopreserved at passages 5 and 10, with full characterisation at passages 10–12. Two cell lines (BD001 and CT003) were reprogrammed in Canada at Prof. Andreazza’s lab (University of Toronto) using the same method, then expanded, characterised, and stored in Australia alongside the other cell lines.

### Germ layer differentiation

iPSCs were differentiated into ectoderm, mesoderm, and endoderm using the STEMDiff™ Trilineage Differentiation Kit (StemCell Technologies, Canada). Cells were plated on Matrigel-coated plates (Corning, NY, US) in mesoderm/endoderm or ectoderm plating media. After 24 hours, the appropriate STEMdiff™ Trilineage medium was added and changed daily until day 5 (mesoderm and endoderm) or day 7 (ectoderm). At these time points, cells were passaged and cultured for two additional days for immunofluorescence analysis and RNA harvest.

### Karyotyping

iPSCs were harvested with TrypLE (Gibco, Thermo Fisher Scientific, MA, US), and the pellets were stored at −20°C until analysis. Molecular karyotypes were assessed using the Illumina Infinium GSA-24 v3.0 array with the human reference sequence GRCh38/hg38 (Dec 2013) (VCGS, Melbourne, Australia) (Supplementary file 1).

### Mycoplasma testing

iPSCs were confirmed to be mycoplasma-free using the MycoAlert® Mycoplasma Detection Kit (Lonza Bioscience, Switzerland) per the manufacturer’s instructions. iPSCs were cultured in T25 flasks with 8 ml of media, and 1 ml of media was collected after 48 hours for testing. Results were interpreted against assay controls: values <0.9 were negative, and >1.2 were positive for mycoplasma (Supplementary Table 1).

### Flow cytometry

iPSCs were dissociated into single cells with TrypLE Select for 3 minutes at 37°C. The cell suspension was stained with BD Horizon™ Fixable Viability Stain 570 (BD Biosciences, NJ, US) for 15 minutes, fixed with 4% PFA for 15 minutes, and incubated with conjugated antibodies for iPSC markers CD9, SSEA4, and EPCAM (Supplementary Table 2) for 30 minutes. Samples were analysed using a BD Canto flow cytometer (BD Biosciences, NJ, US), with results shown in histograms (Supplementary file 3).

### Immunocytochemistry

Cells were cultured until approximately 60% confluency, then washed in PBS and fixed in 4% paraformaldehyde for 15 min at room temperature. Permeabilization solution (0.5% Triton X-100 in PBS) was added for 15 min then blocked in blocking solution (3% BSA in PBS) for 1 hour at room temperature. Appropriate primary antibodies diluted in blocking solution was added (Supplementary Table 2) and incubated for 3 hours at room temperature. Cells were washed three times in PBS, and an appropriate secondary antibody (Supplementary Table 2) diluted in blocking buffer was added and incubated at room temperature for 1 hour. Cells were washed a further three times with PBS, with NucBlue (Invitogen, Thermo Fisher Scientific, MA, US) cell stain (Hoechst 33342) added into the last wash and incubated for five minutes at room temperature before imaging. Cells were imaged using the EVOS M700 Imaging System (Invitrogen, Thermo Fisher Scientific, MA, US). Images are presented unprocessed.

### Proliferation

iPSCs were seeded into 24-well plates coated with Geltrex and maintained for 72 hours in mTeSR Plus medium with daily media changes. Cells were washed with PBS, and then gDNA was extracted using QuickExtract DNA Extraction Solution 1.0 (Biosearch Technologies, Middlesex, UK) as per manufacturer instructions. Total gDNA was measured using the NanoDrop (Thermo Fisher) at 0h, 24h, 48h, and 72 h. Analysis shows normalised values to 0h.

### RNA extraction and qPCR

RNA was extracted using RNeasy® Mini Kits (Qiagen, Germany) and reverse-transcribed into cDNA with the Maxima H Minus First Strand cDNA Synthesis Kit (Thermo Fisher Scientific, MA, US). cDNA concentration was quantified using the Quant-iT™ OliGreen® ssDNA Assay Kit (Thermo Fisher Scientific, MA, US). Gene expression was measured via real-time quantitative PCR (qPCR) on a QuantStudio 3 system (Thermo Fisher Scientific, MA, US) following this protocol: 95°C for 7 min, 40 cycles of 95°C for 30s and 60°C for 1 min (with data acquisition), 60°C for 30s, 55–95°C (with data acquisition), and 20°C for 10s. Gene expression was quantified using the ΔΔCt method and normalized to cDNA concentration. Primers sequences are in Supplementary Table 2.

### Data analysis

For proliferation and qPCR, all analyses were conducted using GraphPad Prism version 10.3.1. All parameters of gene expression and proliferation assays were recorded and grouped as the BD or control group, with 12 input measures for each group. The Shapiro-Wilk test was used to test for normality of data distribution, and multiple T-tests were used for the analysis.

### Cortical network differentiation

For neural progenitor differentiation, iPSCs were seeded at 2 × 10^6^ cells per well in Matrigel-coated 6-well plates using STEMDiff Neural Induction Media with SMADi supplements (StemCell Technologies, Canada). Media changes were performed daily, and cells were passaged every 7 days until passage 3. At passage 3, cells were either expanded for cryopreservation using STEMDiff Neural Progenitor Medium (StemCell Technologies, Canada) or seeded at 4 × 10^4^ cells per well in PLO/Laminin-coated 24-well plates for ‘cortical network’ (a mix of neurons and astrocytes - CNs) differentiation. The cortical network protocol lasted 35 days, with media changes every other day using the BrainPhys hPSC Neuron Kit (StemCell Technologies, Canada). Neural progenitor cells and cortical networks were characterised via immunocytochemistry and gene expression analysis as described above.

### RNA sequencing

CNs cells were harvested using Trizol for RNA extraction and whole transcriptome sequencing. The libraries were prepared using Illumina Total RNA with Ribo-Zero Plus library prep kit and sequenced on Illumina NovaSeq X Sequencing platform with 150bp PE read length, 100M reads per sample for whole transcriptome sequencing. Paired-end raw read data was processed using the Galaxy platform^9^ (Galaxy | Australia (usegalaxy.org.au), accessed September 2024). Filtered reads were aligned to the human reference genome (GRCh38) using HISAT2 in RF-stranded mode^10^. Alignments were filtered based on quality, retaining only reads with a Phred score of ≥10 in the BAM output. Gene-level read counts were quantified with featureCounts ^11^. The quantified read counts for individual samples were combined into an m × n matrix for differential expression analysis. Normalization was performed using the median of ratios method in the DESeq2 package in R^12^. Genes with low expression (<1 CPM in the smallest group size) were excluded from further analysis. Differential gene expression analysis was conducted using DESeq2, with an adjusted p-value threshold of <0.05 to identify significantly differentially expressed genes. The Benjamini-Hochberg method was applied to adjust for multiple comparisons. To account for potential surrogate variables not captured by recorded covariates, the sva package in R^13^ was used applying the formula: Diagnosis + Sample Type.

### Gene set enrichment analysis (GSEA)

Gene set enrichment analysis was conducted using the clusterProfiler package in R^14^ to identify enriched biological pathways, with gene lists pre-ranked by multiplying the sign of log fold changes and the log10-transformed p-values from differential analysis. The KEGG database was used to explore molecular pathways, with significant pathways identified based on an adjusted p-value of <0.05.

## Results

### iPSC line characterisation

All 24 lines presented with the expected iPSC morphology – tightly packed colonies with a high nucleus-to-cytoplasm ratio and pronounced nucleoli (Figure 1A). Expression of pluripotency markers was verified by immunofluorescence (TRA-1-60 and OCT4 – Figure 1B) and flow cytometry (SSEA4, EPCAM and CD9 – Figure 1D). The differentiation potential of the cell lines was demonstrated by the positive expression of ectoderm (Nestin, PAX6 and β-tubulin), endoderm (AFP, SOX17 and FOXA2) and mesoderm (NCAM, Brachyury T and SMA) markers after five to seven days of differentiation by immunofluorescence and qPCR (Figure 1C and 1F). The absence of mycoplasma contamination was confirmed by luminescence assay (Supplementary Table 1). In addition, all cell lines presented with normal molecular karyotype and identical genotype corresponding to the PBMCs when analysed by SNP array (Supplementary File 1). No significant differences were observed between BD and control cell lines regarding the expression of marker genes or proliferation potential (Figure 1E and 1G). BD001 and CT003 lines are also characterised and used to generate cerebral organoids within Prof Andreazza’s group^15^.

**Figure 1.**
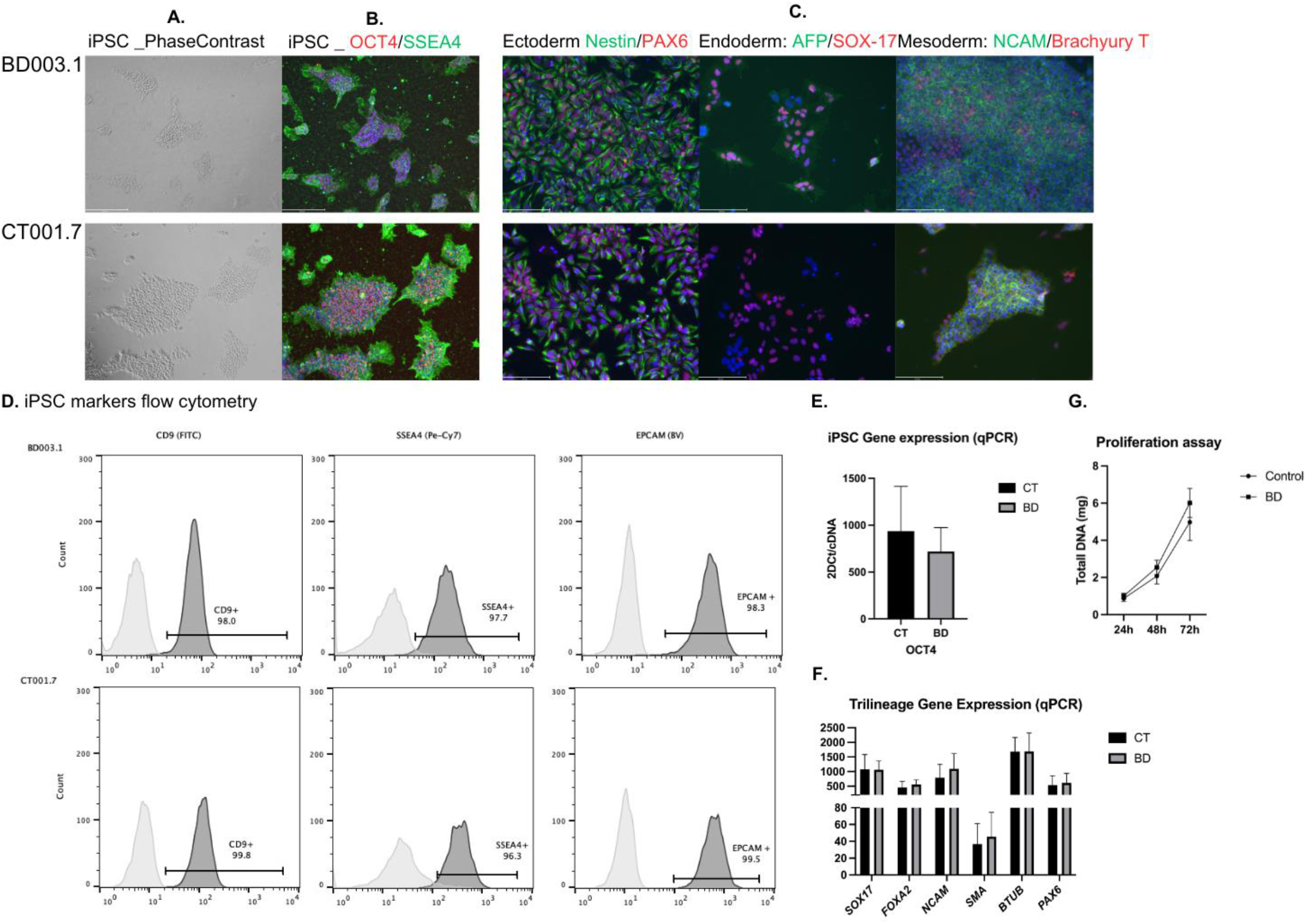
iPSC characterisation. Legend: All 20 iPSC lines exhibited typical morphology (Figure 1A), expressed pluripotency markers by immunofluorescence (TRA-1-60, OCT4; Figure 1B) and flow cytometry (SSEA4, EPCAM, CD9; Figure 1D), and demonstrated differentiation potential into ectoderm, mesoderm, and endoderm lineages by immunofluorescence and qPCR (Figure 1C, 1F). No significant differences were observed between BD and control lines in key gene expression or proliferation (Figure 1E, 1G)—additional information for all lines in supplementary materials. Panels A, B, C and D are representative; full data for all cell lines are in supplementary files 3 to 6.

### CNs

On day 35, all lines expressed markers for mature neurons and astrocytes, as verified by immunofluorescence (MAP2 = neurons and GFAP = astrocytes) and qPCR (β-tubulin = neurons and S100B = astrocytes) (Figure 2).

**Figure 2.**
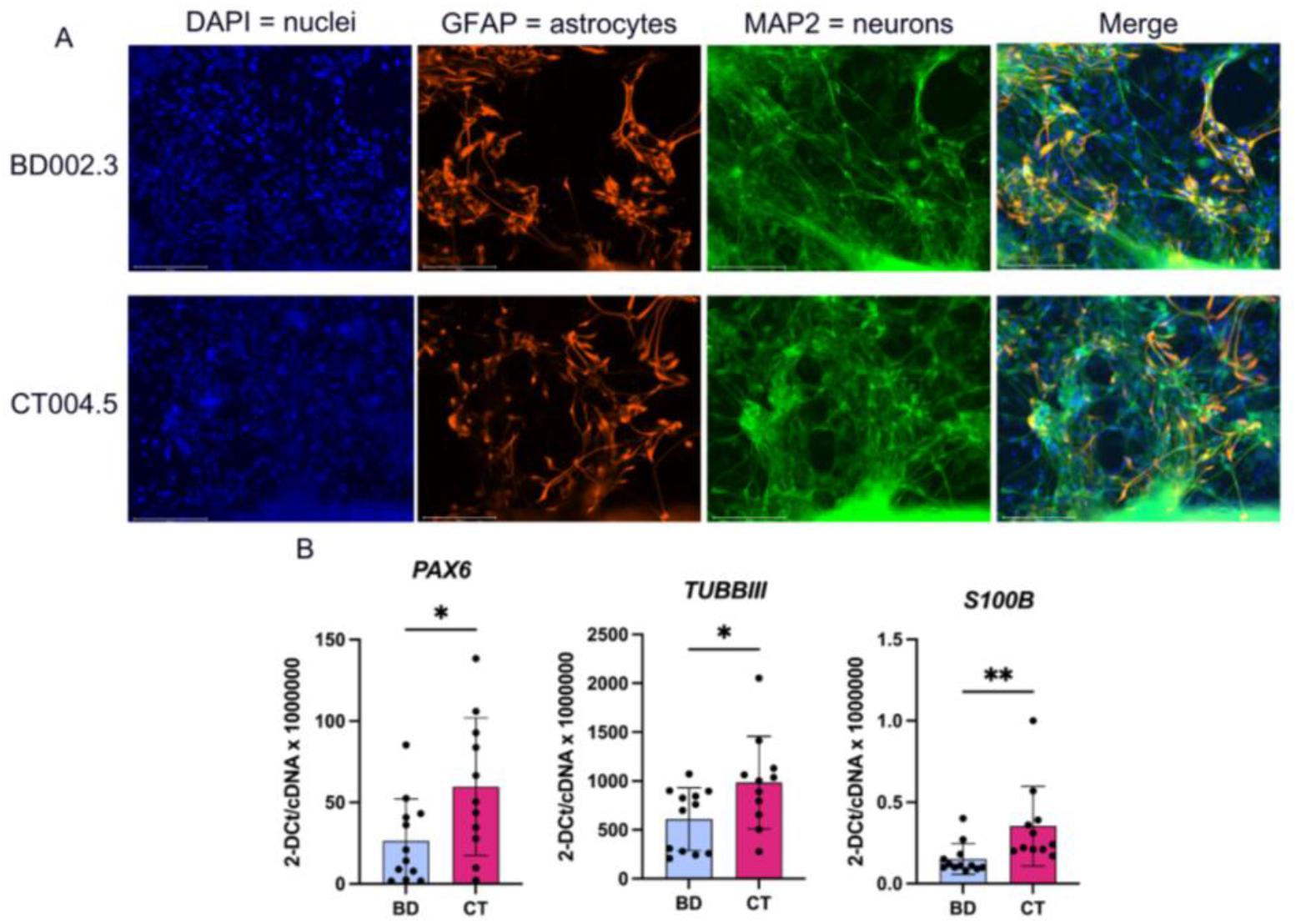
CNs characterisation. Legend: A: Representative image of 2 cell lines neuronal and astrocytic markers. Immunofluorescence: blue, DAPI = Nuclei; green, MAP2 = neurons; orange, GFAP = astrocytes; merge. Scale bar = 150μm. B: qPCR results for neuroprogenitor, neurons and astrocytes. Data shows as mean in SD from the paired t-test for PAX6 (p= 0.031) and β-tubulin (p= 0.035), and Mann-Whitney test for S100B (p= 0.001).

### RNA Seq

CNs whole transcriptome sequencing yielded a total of 3.5 billion paired-end reads, ~18 million reads per sample, with a minimum of 4.6 GBases of data per sample. Differential gene expression analysis identified 1457 genes differentially expressed in BD; of these, 910 genes were downregulated. After correction for multiple testing 3 genes remained differentially expressed (adj. p <0.05) between disease and control samples (2 downregulated: *LYSMD3* and *TINCR*; 1 upregulated: *ENSG00000281181*).

### GSEA

GSEA identified 191 enriched pathways, of which 171 were downregulated in bipolar disorder cells and 20 were upregulated. The top 10 pathways (q-values and NES) are shown in Table 2 (the complete list is in the supplementary table 3).

**Table 2.**
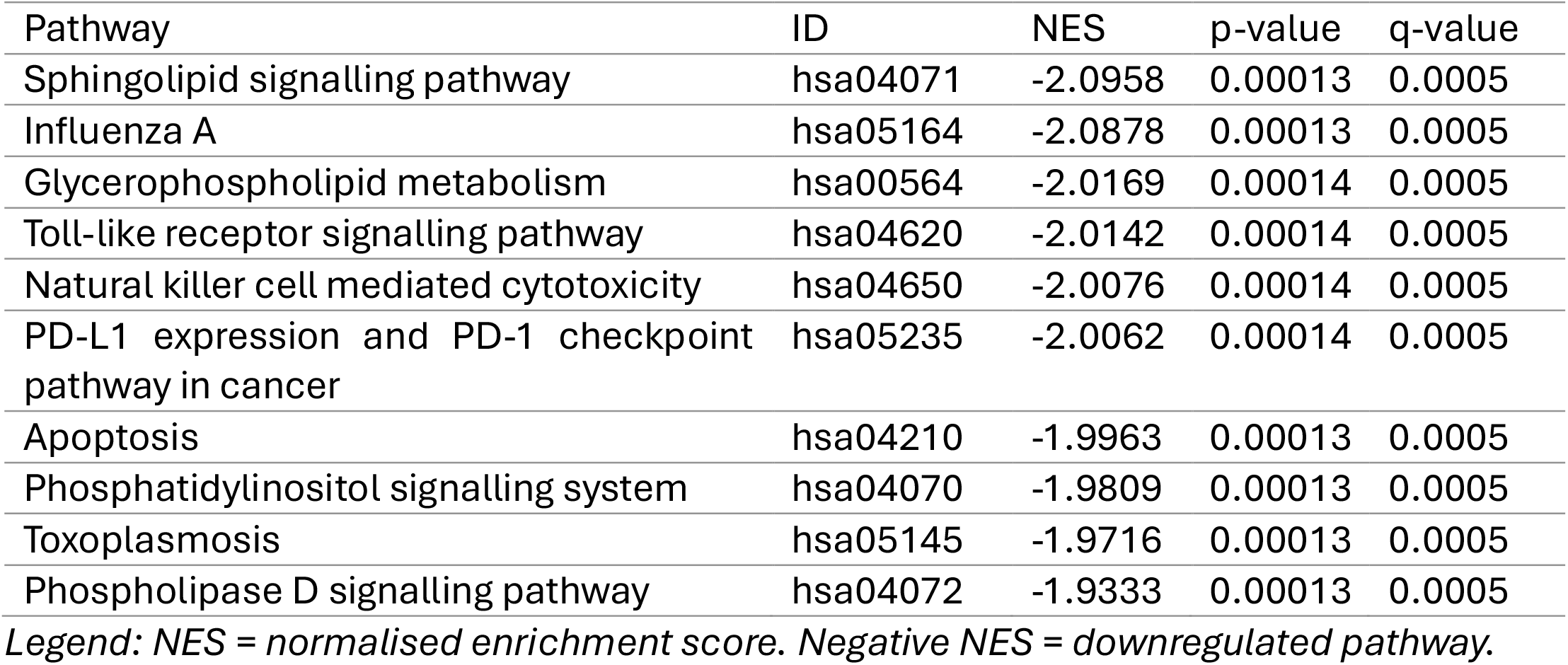
GSEA top ten pathways. Legend: NES = normalised enrichment score. Negative NES = downregulated pathway.

### Toll-like receptors signalling pathway

Top ten pathways identified by GSEA were all downregulations in BD relative to controls. We chose to focus on the Toll-Like Receptor (TLR) pathway for the deeper analysis due to its well-established role in neuroinflammation and immune signalling and its potential role in BD^16^. Investigating TLR dysfunction in BD may provide more immediate mechanistic insights into its inflammatory underpinnings, complementing current knowledge.

Figure 3A shows the top genes affecting the enrichment score for the TLR signalling pathway. Of these, several were selected for confirmation using qPCR (Figure 3B). Among top contributors for the enrichment of this pathway in our dataset, gene expression of 2 human TLRs was confirmed, i.e., TLR3, TLR4.

**Figure 3.**
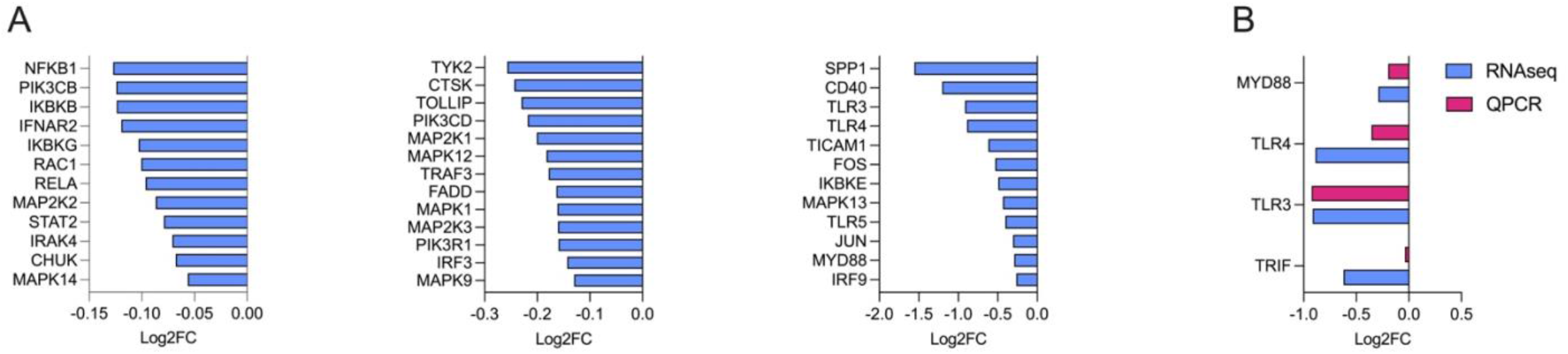
TLR Pathway genes whole transcriptome sequence and qPCR. Legend: A. Log2 fold change gene expression levels of main genes in the TLR signalling pathway. B. Gene expression confirmation by qPCR versus whole transcriptome sequence. MYD88 = MYD88 Innate Immune Signal Transduction Adaptor; TLR4 = Toll-Like Receptor 4; TLR3 = Toll-Like Receptor 3; TRIF or TICAM1= TIR Domain Containing Adaptor Molecule 1.

## Discussion

Using episomal vectors, we generated 12 iPSC lines from PBMCs isolated from adult participants diagnosed with BD; these lines were fully characterised which allows further investigation of the underlying mechanisms of BD and the identification of new treatments and treatment targets. For this project, we have differentiated the iPSCs into cortical networks. Our model combines processes and durations that recapitulate the initial stages of human foetal cortical development: from iPSCs to cortical stem and progenitor cells, followed by a period of cortical neurogenesis and late astrocyte biogenesis, permitting neuronal and astrocyte cell differentiation and maturation to acquire network formation (Figure 2D and Supplementary file 6).

Whole transcriptome sequencing and analysis identified three differentially expressed genes between cells from participants with BD and participants without a history of mental health disorders: *LYSMD3* is a protein-coding gene with a peptidoglycan binding domain involved in Golgi organisation and the transcriptional stress response^17^; *TINCR* is a long-non-coding RNA (lncRNA) associated with cancer and cancer progression^18^; *ENSG00000281181* is a novel lncRNA similar to YY1 associated with myogenesis^19^. To the best of these authors’ knowledge, none of these genes has been previously associated with BD.

Of the 191 pathways that were significantly enriched in our data set, at least 10% are associated with the immune system, infectious and immune diseases. This hints that a profound immune dysregulation is associated with BD, especially in the context of the immune system as a signalling network. The Toll-like receptor signalling pathway (TLR) pathways were featured in the top 10 pathways in our analysis. Toll-like receptors (TLR) are pattern recognition receptors that detect both external pathogens (PAMPs) and internal damage signals (DAMPs). When activated, TLRs induce the NF-κβ signalling pathway (also downregulated in our sample), triggering inflammasome activation and cytokine secretion^20^. This is particularly relevant in the context of peripheral immune cells secreting inflammatory cytokines and the chronic low-grade inflammation associated with BD^21^.

Alternatively, genetic studies in BD have associated a higher risk of early onset BD with polymorphisms of the *TLR4* (*rs*1927914 A and *rs*11536891 T, homozygous)^22^ and *TLR2* genes (*rs*3804099 T and *rs*4696480 T, homozygous)^23^. The *TLR4* rs1927914 polymorphism has been suggested to interfere with the promoter activity, causing dysregulation in the receptor expression^24^, where the G nucleotide is predicted to disrupt the C/EBP repressor transcription factor site, resulting in increased TLR4 expression^25^. Similarly, the *TLR4 rs*11536891 polymorphism has been suggested as a regulatory region where the C nucleotide could rescind miRNA binding sites with expected increased expression of TLR4^26^. Thus, the AA and TT genotypes more prevalent in BD would be associated with lower expression of TLR4, corroborating the findings of our study.

Our results do not show the downregulation of TLR2 expression in BD, which may be due to TLR2’s interaction with external factors that may be related to BD. For example, the genotypes associated with BD indicate a “low cytokine-inducer” phenotype because their counterpart *TLR2* rs4696480 TT was associated with higher susceptibility to spontaneous bacterial peritonitis^27^ and the *TLR2* rs3804099 C allele was associated with higher sepsis mortality rate^28^. Moreover, a combined effect of *TLR2* rs3804099 TT genotype and reported childhood sexual abuse on the age of onset in BD has been reported^29^. A modulatory relation is also noted between *Toxoplasma gondii* exposure (seropositivity associated with BD) and the *TLR2* polymorphism^30^. Comparably, *TLR2* and *TLR4* polymorphisms have been associated with increased susceptibility to infectious diseases^31^. The polymorphisms referred to here were identified in a sample of patients (n=572) and controls (n=202) of French descent^22,23^, but no such association was shown in large GWAS studies ^32,33^.

In addition to their immune functions, TLRs contribute to central nervous system plasticity, neuronal growth, circuit development, and cognitive processes^34^. While highly expressed in immune-related cells, TLRs are also expressed in microglia, astrocytes, oligodendrocytes and neurons^35^; thus, recent evidence suggests that TLRs are mediators of neuroplasticity. TLRs have been linked to neural progenitor cells function^34^. In particular, TLR4 deficiency enhances NPCs proliferation, and these are then more likely to differentiate into neurons (instead of astrocytes). Despite increased proliferation and differentiation, TLR4 deficiency results in lower neuronal survival and maturation^36^. In the context of neurodegenerative diseases, it is postulated that TLR4 activation might have a dual action: clearance of toxins and myelination or, in scenarios of excessive activation, the more established effects of cytokine release promoting neuroinflammation and neuronal death. Microglial activation of TLR4 is suggested to increase phagocytic activity, facilitating the clearance of Abeta deposits in Alzheimer’s disease^37^. In astrocytes, TLR4 activation increases the formation of excitatory synapses^38^, while in neurons, it increases the action potential conductance^39^. TLR4 has been shown to support oligodendrocyte progenitor replacement and chronic functional recovery after spinal injury, supporting remyelination^40^. The pleiotropic effects of TLR4 activation have instigated the search for modulators of TLR4 with receptors agonists and antagonists being proposed for different stages of various diseases.

Altogether, our results can be viewed within the neuroprogression hypothesis of BD^41^, suggesting that inherited variations of *TLR4* may be associated with BD. Downregulation or lower expressing variants of *TLR4* influence the reduced hippocampal volume associated with BD due to decreased neuron maturation during developmental periods, which could lead to cognitive impairments in the future. Therefore, we can hypothesise that genetic predisposition to immune dysregulation may increase susceptibility to environmental risk factors, potentially explaining the link between bipolar disorder and early-life exposure to various infections, particularly during crucial periods of neurodevelopment when the brain is highly sensitive to external influences. This parallels the two-hit hypothesis which has been well articulated in schizophrenia^43^. Indeed, dysregulated inflammatory and immune pathways are linked to BD pathophysiology. From infections during the prenatal period – maternal influenza infection and increased risk of BD^44^-through the association of BD with autoimmune disease – thyroiditis^45^ - to the low grade but abnormal activation of inflammatory mediators during mood episodes (mania and depression) in BD^46^.

Our analysis also highlighted other intriguing pathways downregulated in BD. For instance, sphingolipid signalling and glycerophospholipid metabolism pathways were associated with BD through lipidomic studies conducted on post-mortem brain tissue^47^ and blood samples^48^ of patients with bipolar disorder, which identified increased levels of ceramides, a type of sphingolipid. However, these changes were also affected by factors such as age, cardiovascular comorbidities, and medication, emphasising the complexity of their role in BD. Another interesting pathway identified is apoptosis, which is suggested to be associated with BD brain volume loss and cognitive decline^49,50^.

While this study provides valuable insights into the molecular mechanisms underlying bipolar disorder (BD) using iPSC-derived neurons, several limitations must be acknowledged. Although highly relevant for studying patient-specific neuronal phenotypes, the use of iPSC-derived models presents inherent challenges. Two-dimensional (2D) neuronal cultures, while amenable to high-throughput analyses, lack the cellular complexity and network organisation of the human brain. Our cortical networks are a simplified model of the foetal brain, containing cells that primarily come from ectodermal differentiation – neurons, astrocytes and NPCs. Incorporating microglia-containing cultures or advanced co-culture systems may offer a more physiologically relevant environment for studying immune-related pathways, such as TLR signalling. iPSC-based studies are subject to variability arising from donor differences, reprogramming efficiency, and differentiation protocols. While we employed rigorous standardisation and included both BD patients and healthy controls, larger sample sizes and multi-line (more than one clone per participant) comparisons will be necessary to improve the generalizability of our findings. We want to highlight, though, that this is the biggest sample size used in a BD iPSC-derived study. Future investigation is needed to validate and evaluate the potential role of other genes and pathways identified here.

To our knowledge, this study is the first to report TLR pathway disruptions in iPSC-derived neurons and astrocytes from BD patients. Our findings emphasise the cell-type specificity of transcriptomic changes in BD and underscore the value of patient-derived neuronal models in identifying key molecular pathways that may contribute to understanding BD pathophysiology and developing targeted therapies^51^.

## Acknowledgments

The authors would like to extend sincere gratitude to all study participants for their valuable time and sample contributions to this research. Without their willingness to share, participate and improve future outcomes, this study would not have been possible.

This project was supported by CREDIT: The CRE for the Development of Innovative Therapies for Psychiatric Disorders (GNT1153607) and the Baszucki Brain Research Fund and Milken Institute Center for Strategic Philanthropy. MB is supported by a NHMRC Leadership 3 Investigator grant (GNT2017131). JHK is supported by the Australian Research Council Future Fellowship (FT220100351).

## Author Contributions

Conceptualisation: CCB, KW, AA, OD, MB

Methodology:

Recruitment: BP, CCB, GR, AS, BA, RZ

Lab-based experiments: BP, CCB, ZL, ME, BS, CS, DH, AA, HJJ

Data analysis: BP, TT, KW

Writing:

Manuscript preparation: BP, ME, BS, CS

Review and editing: BP, CCB, ZL, ME, BS, CS, TT, DH, GR, AS, BA, RZ, AA, HJJ, OD, JHK, MB, KW

Funding acquisition: BP, CCB, KW, JHK, OD, MB

