## Supplementary files 1-5 for "The toll-like receptor signalling pathway is altered in iPSC-derived cortical networks from people with bipolar disorder"

Victorian Clinical Genetics Services
Murdoch Children's Research Institute
Flemington Road, Parkville, VIC 3052 Australia
P +61 1300 118247 F +61 3 8341 6366
W vcgs.org.au ABN 51 007 032 760

leaders in genetic health

LIMS Report #: 901660

**Patient:** CT001.7 P13 Ct001.7 P13

**DOB:** Unknown
**Sex:** Unknown
**VCGS sample ID:** 22C103778
**Date collected:** 05-May-2022
**Date received:** 05-May-2022
**Date reported:** 20-Jun-2022
**Source:** Cell Pellet
**Ext. sample ID:**
**Ext. patient ID:**

Dr. Katerina Vlahos
Murdoch Children's Research Institute
Level 5/Flemington Road
PARKVILLE VIC 3052

**CC:** lps.gene

### Cytogenetics Laboratory

| | |
|---|---|
| **Clinical details** | Cell line |
| **Specimen source** | Cell Pellet |

#### Molecular karyotype

| | |
|---|---|
| **Array type** | Illumina Infinium GSA-24 v3.0 |
| **Resolution** | 0.50Mb |
| **Assembly** | hg19 / GRCh37 (Feb 2009) |
| **Molecular karyotype** | arr(X,1-22)x2 |
| **Result** | NO ANEUPLOIDIES DETECTED |

**Interpretation**

Female molecular karyotype. No aneuploidies were detected in this sample.

*Molecular karyotyping is limited in its ability to detect low grade mosaicism and genomic copy number changes below the resolution stated. Balanced rearrangements and Robertsonian translocations will not be detected. This test does not exclude single gene disorders caused by sequence mutations or trinucleotide repeat expansions (such as fragile X syndrome, Huntington disease, some spinocerebellar ataxias, Friedreich ataxia and myotonic dystrophy). Testing for fragile X syndrome should be considered in individuals with developmental delay/ intellectual disability. Further genomic based testing may be considered if there remains a high suspicion of a monogenic disorder. Copy number variants that do not contain genes, are well established polymorphisms, or are assessed as being of unlikely clinical significance (based on "ACMG Technical Standards for the Interpretation and Reporting of Constitutional Copy Number Variants"), will not be reported. The classification is based on the current scientific evidence available at the time of reporting. Reporting of regions of homozygosity (>5Mb) is dependent on referral setting and clinical indication. CNVs that contain autosomal recessive genes will not be reported unless there is specific clinical relevance, high carrier population frequency or a history of consanguinity. Please contact the laboratory if there is a family history of a known recessive disorder or a clinically suspected recessive condition. This testing was performed on a standard SNP microarray platform which may not have sufficient probe coverage to detect clinically relevant CNVs related to this patient's specific clinical features. Please contact the laboratory if a higher resolution microarray may be required for a specific gene or genetic condition. This test does not exclude the possibility of tissue limited mosaicism and further testing of an alternative tissue may be considered if clinically indicated. Interpretation is based on the UCSC GRCh37/hg19 human reference sequence.*

Validated: 20-Jun-2022 by David Francis

Enquiries: +61 1300 11 8247

PATHOLOGY REPORT

**END OF TEST REPORT**

**FINAL REPORT**

RCPA The Royal College of Pathologists of Australasia

NATA

Accredited for compliance with NPAAC Standards and ISO 15189

Accreditation number: 3171

Victorian Clinical Genetics Services
Murdoch Children's Research Institute
Flemington Road, Parkville, VIC 3052 Australia
P +61 1300 118247 F +61 3 8341 6366
W vcgs.org.au ABN 51 007 032 760

VCGS leaders in genetic health

LIMS Report #: 901655

**Patient:** CT004.5 P12 Ct004.5 P12

**DOB:** Unknown
**Sex:** Unknown
**VCGS sample ID:** 22C103776
**Date collected:** 05-May-2022
**Date received:** 05-May-2022
**Date reported:** 20-Jun-2022
**Source:** Cell Pellet
**Ext. sample ID:**
**Ext. patient ID:**

Dr. Katerina Vlahos
Murdoch Children's Research Institute
Level 5/Flemington Road
PARKVILLE VIC 3052

**CC:** lps.gene

### Cytogenetics Laboratory

| | |
|---|---|
| **Clinical details** | Cell line |
| **Specimen source** | Cell Pellet |

#### Molecular karyotype

| | |
|---|---|
| **Array type** | Illumina Infinium GSA-24 v3.0 |
| **Resolution** | 0.50Mb |
| **Assembly** | hg19 / GRCh37 (Feb 2009) |
| **Molecular karyotype** | arr(X,1-22)x2 |
| **Result** | NO ANEUPLOIDIES DETECTED |

**Interpretation**

Female molecular karyotype. No aneuploidies were detected in this sample.

*Molecular karyotyping is limited in its ability to detect low grade mosaicism and genomic copy number changes below the resolution stated. Balanced rearrangements and Robertsonian translocations will not be detected. This test does not exclude single gene disorders caused by sequence mutations or trinucleotide repeat expansions (such as fragile X syndrome, Huntington disease, some spinocerebellar ataxias, Friedreich ataxia and myotonic dystrophy). Testing for fragile X syndrome should be considered in individuals with developmental delay/ intellectual disability. Further genomic based testing may be considered if there remains a high suspicion of a monogenic disorder. Copy number variants that do not contain genes, are well established polymorphisms, or are assessed as being of unlikely clinical significance (based on "ACMG Technical Standards for the Interpretation and Reporting of Constitutional Copy Number Variants"), will not be reported. The classification is based on the current scientific evidence available at the time of reporting. Reporting of regions of homozygosity (>5Mb) is dependent on referral setting and clinical indication. CNVs that contain autosomal recessive genes will not be reported unless there is specific clinical relevance, high carrier population frequency or a history of consanguinity. Please contact the laboratory if there is a family history of a known recessive disorder or a clinically suspected recessive condition. This testing was performed on a standard SNP microarray platform which may not have sufficient probe coverage to detect clinically relevant CNVs related to this patient's specific clinical features. Please contact the laboratory if a higher resolution microarray may be required for a specific gene or genetic condition. This test does not exclude the possibility of tissue limited mosaicism and further testing of an alternative tissue may be considered if clinically indicated. Interpretation is based on the UCSC GRCh37/hg19 human reference sequence.*

Validated: 20-Jun-2022 by David Francis

Enquiries: +61 1300 11 8247

**END OF TEST REPORT**

**FINAL REPORT**

PATHOLOGY REPORT

Victorian Clinical Genetics Services
Murdoch Children's Research Institute
Flemington Road, Parkville, VIC 3052 Australia
P +61 1300 118247 F +61 3 8341 6366
W vcgs.org.au ABN 51 007 032 760

VCGS
leaders in genetic health

LIMS Report #: 850334

**Patient:** CT005.1 P10 Ct005.1 P10

**DOB:** Unknown
**Sex:** Unknown
**VCGS sample ID:** 21C111364
**Date collected:** 14-Dec-2021
**Date received:** 14-Dec-2021
**Date reported:** 05-Jan-2022
**Source:** Cell Pellet
**Ext. sample ID:**
**Ext. patient ID:**

Dr. Katerina Vlahos
Murdoch Childrens Research Institute
Level 5/Flemington Road
PARKVILLE VIC 3052

### Cytogenetics Laboratory

| | |
|---|---|
| **Clinical details** | Cell line |
| **Specimen source** | Cell Pellet |

#### Molecular karyotype

| | |
|---|---|
| **Array type** | Illumina Infinium GSA-24 v3.0 |
| **Resolution** | 0.50Mb |
| **Assembly** | hg19 / GRCh37 (Feb 2009) |
| **Molecular karyotype** | arr(X,1-22)x2 |
| **Result** | NO ANEUPLOIDIES DETECTED |

**Interpretation**

Female molecular karyotype. No aneuploidies were detected in this sample.

*Molecular karyotyping is limited in its ability to detect low grade mosaicism and genomic copy number changes below the resolution stated. Balanced rearrangements and Robertsonian translocations will not be detected. This test does not exclude single gene disorders caused by sequence mutations or trinucleotide repeat expansions (such as fragile X syndrome, Huntington disease, some spinocerebellar ataxias, Friedreich ataxia and myotonic dystrophy). Testing for fragile X syndrome should be considered in individuals with developmental delay/ intellectual disability. Further genomic based testing may be considered if there remains a high suspicion of a monogenic disorder. Copy number variants that do not contain genes, are well established polymorphisms, or are assessed as being of unlikely clinical significance (based on “ACMG Technical Standards for the Interpretation and Reporting of Constitutional Copy Number Variants”), will not be reported. The classification is based on the current scientific evidence available at the time of reporting. Reporting of regions of homozygosity (>5Mb) is dependent on referral setting and clinical indication. CNVs that contain autosomal recessive genes will not be reported unless there is specific clinical relevance, high carrier population frequency or a history of consanguinity. Please contact the laboratory if there is a family history of a known recessive disorder or a clinically suspected recessive condition. This testing was performed on a standard SNP microarray platform which may not have sufficient probe coverage to detect clinically relevant CNVs related to this patient’s specific clinical features. Please contact the laboratory if a higher resolution microarray may be required for a specific gene or genetic condition. This test does not exclude the possibility of tissue limited mosaicism and further testing of an alternative tissue may be considered if clinically indicated. Interpretation is based on the UCSC GRCh37/hg19 human reference sequence.*

Validated: 05-Jan-2022 by David Francis

Enquiries: +61 1300 11 8247

PATHOLOGY REPORT

**END OF TEST REPORT**

**FINAL REPORT**

VCGS
leaders in genetic health

Victorian Clinical Genetics Services
Murdoch Children's Research Institute
Flemington Road, Parkville, VIC 3052 Australia
P +61 1300 118247 F +61 3 8341 6366
W vcgs.org.au ABN 51 007 032 760

LIMS Report #: 850335

**Patient:** CT006.10 P10 Ct006.10 P10

**DOB:** Unknown
**Sex:** Unknown
**VCGS sample ID:** 21C111365
**Date collected:** 14-Dec-2021
**Date received:** 14-Dec-2021
**Date reported:** 05-Jan-2022
**Source:** Cell Pellet
**Ext. sample ID:**
**Ext. patient ID:**

Dr. Katerina Vlahos
Murdoch Childrens Research Institute
Level 5/Flemington Road
PARKVILLE VIC 3052

### Cytogenetics Laboratory

| | |
|---|---|
| **Clinical details** | Cell line |
| **Specimen source** | Cell Pellet |

#### Molecular karyotype

| | |
|---|---|
| **Array type** | Illumina Infinium GSA-24 v3.0 |
| **Resolution** | 0.50Mb |
| **Assembly** | hg19 / GRCh37 (Feb 2009) |
| **Molecular karyotype** | arr(X,1-22)x2 |
| **Result** | NO ANEUPLOIDIES DETECTED |

**Interpretation**

Female molecular karyotype. No aneuploidies were detected in this sample.

*Molecular karyotyping is limited in its ability to detect low grade mosaicism and genomic copy number changes below the resolution stated. Balanced rearrangements and Robertsonian translocations will not be detected. This test does not exclude single gene disorders caused by sequence mutations or trinucleotide repeat expansions (such as fragile X syndrome, Huntington disease, some spinocerebellar ataxias, Friedreich ataxia and myotonic dystrophy). Testing for fragile X syndrome should be considered in individuals with developmental delay/ intellectual disability. Further genomic based testing may be considered if there remains a high suspicion of a monogenic disorder. Copy number variants that do not contain genes, are well established polymorphisms, or are assessed as being of unlikely clinical significance (based on "ACMG Technical Standards for the Interpretation and Reporting of Constitutional Copy Number Variants"), will not be reported. The classification is based on the current scientific evidence available at the time of reporting. Reporting of regions of homozygosity (>5Mb) is dependent on referral setting and clinical indication. CNVs that contain autosomal recessive genes will not be reported unless there is specific clinical relevance, high carrier population frequency or a history of consanguinity. Please contact the laboratory if there is a family history of a known recessive disorder or a clinically suspected recessive condition. This testing was performed on a standard SNP microarray platform which may not have sufficient probe coverage to detect clinically relevant CNVs related to this patient's specific clinical features. Please contact the laboratory if a higher resolution microarray may be required for a specific gene or genetic condition. This test does not exclude the possibility of tissue limited mosaicism and further testing of an alternative tissue may be considered if clinically indicated. Interpretation is based on the UCSC GRCh37/hg19 human reference sequence.*

Validated: 05-Jan-2022 by David Francis

Enquiries: +61 1300 11 8247

PATHOLOGY REPORT

**END OF TEST REPORT**

**FINAL REPORT**

Victorian Clinical Genetics Services
Murdoch Children's Research Institute
Flemington Road, Parkville, VIC 3052 Australia
P +61 1300 118247 F +61 3 8341 6366
W vcgs.org.au ABN 51 007 032 760

leaders in genetic health

LIMS Report #: 1011170

**Patient:** #36.8 IPSC P11 #36.8 Ipsc P11

Dr. Katerina Vlahos
Murdoch Children's Research Institute
Level 5/Flemington Road
PARKVILLE VIC 3052

**DOB:** Unknown
**Sex:**
**VCGS sample ID:** 23C105302
**Date collected:** 10-May-2023
**Date received:** 10-May-2023
**Date reported:** 06-Jun-2023
**Source:** Cell Pellet
**Ext. sample ID:**
**Ext. patient ID:**

### Cytogenetics Laboratory

| | |
|---|---|
| **Clinical details** | Cell line |
| **Specimen source** | Cell Pellet |

#### Molecular karyotype

| | |
|---|---|
| **Array type** | Illumina Infinium GSA-24 v3.0 |
| **Resolution** | 0.50Mb |
| **Reference genome** | GRCh38 / hg38 (Dec 2013) |
| **Molecular karyotype** | arr(X,Y)x1,(1-22)x2 |
| **Result** | NO ANEUPLOIDIES DETECTED |

**Interpretation**

Male molecular karyotype. No aneuploidies were detected in this sample.

*Molecular karyotyping is limited in its ability to detect low grade mosaicism and genomic copy number changes below the resolution stated. Balanced rearrangements and Robertsonian translocations will not be detected. This test does not exclude single gene disorders caused by sequence mutations or trinucleotide repeat expansions (such as fragile X syndrome, Huntington disease, some spinocerebellar ataxias, Friedreich ataxia and myotonic dystrophy). Testing for fragile X syndrome should be considered in individuals with developmental delay/ intellectual disability. Further genomic based testing may be considered if there remains a high suspicion of a monogenic disorder. Copy number variants that do not contain genes, are well established polymorphisms, or are assessed as being of unlikely clinical significance (based on "ACMG Technical Standards for the Interpretation and Reporting of Constitutional Copy Number Variants"), will not be reported. The classification is based on the current scientific evidence available at the time of reporting. Reporting of regions of homozygosity (>5Mb) is dependent on referral setting and clinical indication. CNVs that contain autosomal recessive genes will not be reported unless there is specific clinical relevance, high carrier population frequency or a history of consanguinity. Please contact the laboratory if there is a family history of a known recessive disorder or a clinically suspected recessive condition. CNVs involving moderate to high risk cancer susceptibility genes may be reported. However, incidentally ascertained CNVs involving genes assessed as having low to moderate cancer risk will not generally be reported unless they are part of a larger CNV. This testing was performed on a standard SNP microarray platform which may not have sufficient probe coverage to detect clinically relevant CNVs related to this patient's specific clinical features. Please contact the laboratory if a higher resolution microarray may be required for a specific gene or genetic condition. This test does not exclude the possibility of tissue limited mosaicism and further testing of an alternative tissue may be considered if clinically indicated. Interpretation is based on the UCSC GRCh38/hg38 human reference sequence.*

Validated: 06-Jun-2023 by David Francis

Enquiries: +61 1300 11 8247

PATHOLOGY REPORT

**END OF TEST REPORT**

**FINAL REPORT**

Accredited for compliance with NPAAC Standards and ISO 15189

Accreditation number: 3171

Victorian Clinical Genetics Services
Murdoch Children's Research Institute
Flemington Road, Parkville, VIC 3052 Australia
P +61 1300 118247 F +61 3 8341 6366
W vcgs.org.au ABN 51 007 032 760

leaders in genetic health

LIMS Report #: 850331

**Patient:** CT003 P12 Ct003 P12

**DOB:** Unknown
**Sex:** Unknown
**VCGS sample ID:** 21C111362

Dr. Katerina Vlahos
Murdoch Childrens Research Institute
Level 5/Flemington Road
PARKVILLE VIC 3052

**Date collected:** 14-Dec-2021
**Date received:** 14-Dec-2021
**Date reported:** 05-Jan-2022
**Source:** Cell Pellet
**Ext. sample ID:**
**Ext. patient ID:**

### Cytogenetics Laboratory

| | |
|---|---|
| **Clinical details** | Cell line |
| **Specimen source** | Cell Pellet |

#### Molecular karyotype

| | |
|---|---|
| **Array type** | Illumina Infinium GSA-24 v3.0 |
| **Resolution** | 0.50Mb |
| **Assembly** | hg19 / GRCh37 (Feb 2009) |
| **Molecular karyotype** | arr(X,1-22)x2 |
| **Result** | NO ANEUPLOIDIES DETECTED |

**Interpretation**

Female molecular karyotype. No aneuploidies were detected in this sample.

*Molecular karyotyping is limited in its ability to detect low grade mosaicism and genomic copy number changes below the resolution stated. Balanced rearrangements and Robertsonian translocations will not be detected. This test does not exclude single gene disorders caused by sequence mutations or trinucleotide repeat expansions (such as fragile X syndrome, Huntington disease, some spinocerebellar ataxias, Friedreich ataxia and myotonic dystrophy). Testing for fragile X syndrome should be considered in individuals with developmental delay/ intellectual disability. Further genomic based testing may be considered if there remains a high suspicion of a monogenic disorder. Copy number variants that do not contain genes, are well established polymorphisms, or are assessed as being of unlikely clinical significance (based on "ACMG Technical Standards for the Interpretation and Reporting of Constitutional Copy Number Variants"), will not be reported. The classification is based on the current scientific evidence available at the time of reporting. Reporting of regions of homozygosity (>5Mb) is dependent on referral setting and clinical indication. CNVs that contain autosomal recessive genes will not be reported unless there is specific clinical relevance, high carrier population frequency or a history of consanguinity. Please contact the laboratory if there is a family history of a known recessive disorder or a clinically suspected recessive condition. This testing was performed on a standard SNP microarray platform which may not have sufficient probe coverage to detect clinically relevant CNVs related to this patient's specific clinical features. Please contact the laboratory if a higher resolution microarray may be required for a specific gene or genetic condition. This test does not exclude the possibility of tissue limited mosaicism and further testing of an alternative tissue may be considered if clinically indicated. Interpretation is based on the UCSC GRCh37/hg19 human reference sequence.*

Validated: 05-Jan-2022 by David Francis

Enquiries: +61 1300 11 8247

PATHOLOGY REPORT

**END OF TEST REPORT**

**FINAL REPORT**

Victorian Clinical Genetics Services
Murdoch Children's Research Institute
Flemington Road, Parkville, VIC 3052 Australia
P +61 1300 118247 F +61 3 8341 6366
W vcgs.org.au ABN 51 007 032 760

LIMS Report #: 850336

**Patient:** CT007.5 P10 Ct007.5 P10

**DOB:** Unknown
**Sex:** Unknown
**VCGS sample ID:** 21C111366
**Date collected:** 14-Dec-2021
**Date received:** 14-Dec-2021
**Date reported:** 05-Jan-2022
**Source:** Cell Pellet
**Ext. sample ID:**
**Ext. patient ID:**

Dr. Katerina Vlahos
Murdoch Childrens Research Institute
Level 5/Flemington Road
PARKVILLE VIC 3052

### Cytogenetics Laboratory

| | |
|---|---|
| **Clinical details** | Cell line |
| **Specimen source** | Cell Pellet |

#### Molecular karyotype

| | |
|---|---|
| **Array type** | Illumina Infinium GSA-24 v3.0 |
| **Resolution** | 0.50Mb |
| **Assembly** | hg19 / GRCh37 (Feb 2009) |
| **Molecular karyotype** | arr(X,1-22)x2 |
| **Result** | NO ANEUPLOIDIES DETECTED |

**Interpretation**

Female molecular karyotype. No aneuploidies were detected in this sample.

*Molecular karyotyping is limited in its ability to detect low grade mosaicism and genomic copy number changes below the resolution stated. Balanced rearrangements and Robertsonian translocations will not be detected. This test does not exclude single gene disorders caused by sequence mutations or trinucleotide repeat expansions (such as fragile X syndrome, Huntington disease, some spinocerebellar ataxias, Friedreich ataxia and myotonic dystrophy). Testing for fragile X syndrome should be considered in individuals with developmental delay/ intellectual disability. Further genomic based testing may be considered if there remains a high suspicion of a monogenic disorder. Copy number variants that do not contain genes, are well established polymorphisms, or are assessed as being of unlikely clinical significance (based on “ACMG Technical Standards for the Interpretation and Reporting of Constitutional Copy Number Variants”), will not be reported. The classification is based on the current scientific evidence available at the time of reporting. Reporting of regions of homozygosity (>5Mb) is dependent on referral setting and clinical indication. CNVs that contain autosomal recessive genes will not be reported unless there is specific clinical relevance, high carrier population frequency or a history of consanguinity. Please contact the laboratory if there is a family history of a known recessive disorder or a clinically suspected recessive condition. This testing was performed on a standard SNP microarray platform which may not have sufficient probe coverage to detect clinically relevant CNVs related to this patient’s specific clinical features. Please contact the laboratory if a higher resolution microarray may be required for a specific gene or genetic condition. This test does not exclude the possibility of tissue limited mosaicism and further testing of an alternative tissue may be considered if clinically indicated. Interpretation is based on the UCSC GRCh37/hg19 human reference sequence.*

Validated: 05-Jan-2022 by David Francis

Enquiries: +61 1300 11 8247

PATHOLOGY REPORT

**END OF TEST REPORT**

**FINAL REPORT**

Victorian Clinical Genetics Services
Murdoch Children's Research Institute
Flemington Road, Parkville, VIC 3052 Australia
P +61 1300 118247 F +61 3 8341 6366
W vcgs.org.au ABN 51 007 032 760

LIMS Report #: 850329

**Patient:** ATCC 5.1 P10 Atcc 5.1 P10

Dr. Katerina Vlahos
Murdoch Childrens Research Institute
Level 5/Flemington Road
PARKVILLE VIC 3052

**DOB:** Unknown
**Sex:** Unknown
**VCGS sample ID:** 21C111360
**Date collected:** 14-Dec-2021
**Date received:** 14-Dec-2021
**Date reported:** 05-Jan-2022
**Source:** Cell Pellet
**Ext. sample ID:**
**Ext. patient ID:**

### Cytogenetics Laboratory

| | |
|---|---|
| **Clinical details** | Cell line |
| **Specimen source** | Cell Pellet |

#### Molecular karyotype

| | |
|---|---|
| **Array type** | Illumina Infinium GSA-24 v3.0 |
| **Resolution** | 0.50Mb |
| **Assembly** | hg19 / GRCh37 (Feb 2009) |
| **Molecular karyotype** | arr(X,1-22)x2 |
| **Result** | NO ANEUPLOIDIES DETECTED |

**Interpretation**

Female molecular karyotype. No aneuploidies were detected in this sample.

*Molecular karyotyping is limited in its ability to detect low grade mosaicism and genomic copy number changes below the resolution stated. Balanced rearrangements and Robertsonian translocations will not be detected. This test does not exclude single gene disorders caused by sequence mutations or trinucleotide repeat expansions (such as fragile X syndrome, Huntington disease, some spinocerebellar ataxias, Friedreich ataxia and myotonic dystrophy). Testing for fragile X syndrome should be considered in individuals with developmental delay/ intellectual disability. Further genomic based testing may be considered if there remains a high suspicion of a monogenic disorder. Copy number variants that do not contain genes, are well established polymorphisms, or are assessed as being of unlikely clinical significance (based on "ACMG Technical Standards for the Interpretation and Reporting of Constitutional Copy Number Variants"), will not be reported. The classification is based on the current scientific evidence available at the time of reporting. Reporting of regions of homozygosity (>5Mb) is dependent on referral setting and clinical indication. CNVs that contain autosomal recessive genes will not be reported unless there is specific clinical relevance, high carrier population frequency or a history of consanguinity. Please contact the laboratory if there is a family history of a known recessive disorder or a clinically suspected recessive condition. This testing was performed on a standard SNP microarray platform which may not have sufficient probe coverage to detect clinically relevant CNVs related to this patient's specific clinical features. Please contact the laboratory if a higher resolution microarray may be required for a specific gene or genetic condition. This test does not exclude the possibility of tissue limited mosaicism and further testing of an alternative tissue may be considered if clinically indicated. Interpretation is based on the UCSC GRCh37/hg19 human reference sequence.*

Validated: 05-Jan-2022 by David Francis

Enquiries: +61 1300 11 8247

PATHOLOGY REPORT

**END OF TEST REPORT**

**FINAL REPORT**

Accredited for compliance with NPAAC Standards and ISO 15189
Accreditation number: 3171

Victorian Clinical Genetics Services
Murdoch Children's Research Institute
Flemington Road, Parkville, VIC 3052 Australia
P +61 1300 118247 F +61 3 8341 6366
W vcgs.org.au ABN 51 007 032 760

LIMS Report #: 1061935

**Patient:** EOI #184.6 P11 Eoi #184.6 P11

Dr. Katerina Vlahos
Murdoch Children's Research Institute
Level 5/Flemington Road
PARKVILLE VIC 3052

**DOB:** Unknown
**Sex:** Unknown
**VCGS sample ID:** 23C112728
**Date collected:** 19-Oct-2023
**Date received:** 19-Oct-2023
**Date reported:** 10-Nov-2023
**Source:** Cell Pellet
**Ext. sample ID:**
**Ext. patient ID:**

**CC:** Ips.gene

### Cytogenetics Laboratory

PATHOLOGY REPORT

| | |
|---|---|
| **Clinical details** | Cell line |
| **Specimen source** | Cell Pellet |

#### Molecular karyotype

| | |
|---|---|
| **Array type** | Illumina Infinium GSA-24 v3.0 |
| **Resolution** | 0.50Mb |
| **Reference genome** | GRCh38 / hg38 (Dec 2013) |
| **Molecular karyotype** | arr(X,1-22)x2 |
| **Result** | NO ANEUPLOIDIES DETECTED |

**Interpretation**

Female molecular karyotype. No aneuploidies were detected in this sample.

*Molecular karyotyping is limited in its ability to detect low grade mosaicism and genomic copy number changes below the resolution stated. Balanced rearrangements and Robertsonian translocations will not be detected. This test does not exclude single gene disorders caused by sequence mutations or trinucleotide repeat expansions (such as fragile X syndrome, Huntington disease, some spinocerebellar ataxias, Friedreich ataxia and myotonic dystrophy). Testing for fragile X syndrome should be considered in individuals with developmental delay/ intellectual disability. Further genomic based testing may be considered if there remains a high suspicion of a monogenic disorder. Copy number variants that do not contain genes, are well established polymorphisms, or are assessed as being of unlikely clinical significance (based on "ACMG Technical Standards for the Interpretation and Reporting of Constitutional Copy Number Variants"), will not be reported. The classification is based on the current scientific evidence available at the time of reporting. Reporting of regions of homozygosity (>5Mb) is dependent on referral setting and clinical indication. CNVs that contain autosomal recessive genes will not be reported unless there is specific clinical relevance, high carrier population frequency or a history of consanguinity. Please contact the laboratory if there is a family history of a known recessive disorder or a clinically suspected recessive condition. This testing was performed on a standard SNP microarray platform which may not have sufficient probe coverage to detect clinically relevant CNVs related to this patient's specific clinical features. Please contact the laboratory if a higher resolution microarray may be required for a specific gene or genetic condition. This test does not exclude the possibility of tissue limited mosaicism and further testing of an alternative tissue may be considered if clinically indicated. Interpretation is based on the UCSC GRCh38/hg38 human reference sequence.*

Validated: 10-Nov-2023 by Vida Petrovic

Enquiries: +61 1300 11 8247

**END OF TEST REPORT**

**FINAL REPORT**

Victorian Clinical Genetics Services
Murdoch Children's Research Institute
Flemington Road, Parkville, VIC 3052 Australia
P +61 1300 118247 F +61 3 8341 6366
W vcgs.org.au ABN 51 007 032 760

VCGS leaders in genetic health

LIMS Report #: 961285

**Patient:** EO1 #87.4 P11 Eo1 #87.4 P11

**DOB:** Unknown
**Sex:** Unknown
**VCGS sample ID:** 22C111969
**Date collected:** 14-Dec-2022
**Date received:** 14-Dec-2022
**Date reported:** 29-Dec-2022
**Source:** Cell Pellet
**Ext. sample ID:**
**Ext. patient ID:**

Dr. Katerina Vlahos
Murdoch Children's Research Institute
Level 5/Flemington Road
PARKVILLE VIC 3052

**CC:** Ips.gene

### Cytogenetics Laboratory

| | |
|---|---|
| **Clinical details** | Cell line |
| **Specimen source** | Cell Pellet |

#### Molecular karyotype

| | |
|---|---|
| **Array type** | Illumina Infinium GSA-24 v3.0 |
| **Resolution** | 0.50Mb |
| **Assembly** | hg19 / GRCh37 (Feb 2009) |
| **Molecular karyotype** | arr(X,1-22)x2 |
| **Result** | NO ANEUPLOIDIES DETECTED |

**Interpretation**

Female molecular karyotype. No aneuploidies were detected in this sample.

*Molecular karyotyping is limited in its ability to detect low grade mosaicism and genomic copy number changes below the resolution stated. Balanced rearrangements and Robertsonian translocations will not be detected. This test does not exclude single gene disorders caused by sequence mutations or trinucleotide repeat expansions (such as fragile X syndrome, Huntington disease, some spinocerebellar ataxias, Friedreich ataxia and myotonic dystrophy). Testing for fragile X syndrome should be considered in individuals with developmental delay/ intellectual disability. Further genomic based testing may be considered if there remains a high suspicion of a monogenic disorder. Copy number variants that do not contain genes, are well established polymorphisms, or are assessed as being of unlikely clinical significance (based on "ACMG Technical Standards for the Interpretation and Reporting of Constitutional Copy Number Variants"), will not be reported. The classification is based on the current scientific evidence available at the time of reporting. Reporting of regions of homozygosity (>5Mb) is dependent on referral setting and clinical indication. CNVs that contain autosomal recessive genes will not be reported unless there is specific clinical relevance, high carrier population frequency or a history of consanguinity. Please contact the laboratory if there is a family history of a known recessive disorder or a clinically suspected recessive condition. This testing was performed on a standard SNP microarray platform which may not have sufficient probe coverage to detect clinically relevant CNVs related to this patient's specific clinical features. Please contact the laboratory if a higher resolution microarray may be required for a specific gene or genetic condition. This test does not exclude the possibility of tissue limited mosaicism and further testing of an alternative tissue may be considered if clinically indicated. Interpretation is based on the UCSC GRCh37/hg19 human reference sequence.*

Validated: 29-Dec-2022 by David Francis

Enquiries: +61 1300 11 8247

PATHOLOGY REPORT

**END OF TEST REPORT**

**FINAL REPORT**

VCGS
leaders in genetic health

Victorian Clinical Genetics Services
Murdoch Children's Research Institute
Flemington Road, Parkville, VIC 3052 Australia
P +61 1300 118247 F +61 3 8341 6366
W vcgs.org.au ABN 51 007 032 760

LIMS Report #: 1061946

**Patient:** EOI #110.1 P11 Eoi #110.1 P11

**DOB:** Unknown
**Sex:** Unknown
**VCGS sample ID:** 23C112733
**Date collected:** 19-Oct-2023
**Date received:** 19-Oct-2023
**Date reported:** 10-Nov-2023
**Source:** Cell Pellet
**Ext. sample ID:**
**Ext. patient ID:**

Dr. Katerina Vlahos
Murdoch Children's Research Institute
Level 5/Flemington Road
PARKVILLE VIC 3052

**CC:** Ips.gene

PATHOLOGY REPORT

### Cytogenetics Laboratory

| | |
|---|---|
| **Clinical details** | Cell line |
| **Specimen source** | Cell Pellet |

#### Molecular karyotype

| | |
|---|---|
| **Array type** | Illumina Infinium GSA-24 v3.0 |
| **Resolution** | 0.50Mb |
| **Reference genome** | GRCh38 / hg38 (Dec 2013) |
| **Molecular karyotype** | arr(X,1-22)x2 |
| **Result** | NO ANEUPLOIDIES DETECTED |

**Interpretation**

Female molecular karyotype. No aneuploidies were detected in this sample.

*Molecular karyotyping is limited in its ability to detect low grade mosaicism and genomic copy number changes below the resolution stated. Balanced rearrangements and Robertsonian translocations will not be detected. This test does not exclude single gene disorders caused by sequence mutations or trinucleotide repeat expansions (such as fragile X syndrome, Huntington disease, some spinocerebellar ataxias, Friedreich ataxia and myotonic dystrophy). Testing for fragile X syndrome should be considered in individuals with developmental delay/ intellectual disability. Further genomic based testing may be considered if there remains a high suspicion of a monogenic disorder. Copy number variants that do not contain genes, are well established polymorphisms, or are assessed as being of unlikely clinical significance (based on "ACMG Technical Standards for the Interpretation and Reporting of Constitutional Copy Number Variants"), will not be reported. The classification is based on the current scientific evidence available at the time of reporting. Reporting of regions of homozygosity (>5Mb) is dependent on referral setting and clinical indication. CNVs that contain autosomal recessive genes will not be reported unless there is specific clinical relevance, high carrier population frequency or a history of consanguinity. Please contact the laboratory if there is a family history of a known recessive disorder or a clinically suspected recessive condition. This testing was performed on a standard SNP microarray platform which may not have sufficient probe coverage to detect clinically relevant CNVs related to this patient's specific clinical features. Please contact the laboratory if a higher resolution microarray may be required for a specific gene or genetic condition. This test does not exclude the possibility of tissue limited mosaicism and further testing of an alternative tissue may be considered if clinically indicated. Interpretation is based on the UCSC GRCh38/hg38 human reference sequence.*

Validated: 10-Nov-2023 by Vida Petrovic

Enquiries: +61 1300 11 8247

**END OF TEST REPORT**

**FINAL REPORT**

RCPA The Royal College of Pathologists of Australasia
NATA WORLD RECOGNISED ACCREDITATION
Accredited for compliance with NPAAC Standards and ISO 15189
ilac-MRA
NATA Accreditation No. 3171

Victorian Clinical Genetics Services
Murdoch Children's Research Institute
Flemington Road, Parkville, VIC 3052 Australia
P +61 1300 118247 F +61 3 8341 6366
W vcgs.org.au ABN 51 007 032 760

VCGS leaders in genetic health

LIMS Report #: 1010836

**Patient:** EOI#98.5 IPSC P11 Eoi#98.5 Ipsc P11

Dr. Katerina Vlahos
Murdoch Children's Research Institute
Level 5/Flemington Road
PARKVILLE VIC 3052

**DOB:** Unknown
**Sex:**
**VCGS sample ID:** 23C105301
**Date collected:** 10-May-2023
**Date received:** 10-May-2023
**Date reported:** 05-Jun-2023
**Source:** Cell Pellet
**Ext. sample ID:**
**Ext. patient ID:**

**CC:** Ips.gene

### Cytogenetics Laboratory

| | |
|---|---|
| **Clinical details** | Cell line |
| **Specimen source** | Cell Pellet |

#### Molecular karyotype

| | |
|---|---|
| **Array type** | Illumina Infinium GSA-24 v3.0 |
| **Resolution** | 0.50Mb |
| **Reference genome** | GRCh38 / hg38 (Dec 2013) |
| **Molecular karyotype** | arr(X,1-22)x2 |
| **Result** | NO ANEUPLOIDIES DETECTED |

**Interpretation**

Female molecular karyotype. No aneuploidies were detected in this sample.

*Molecular karyotyping is limited in its ability to detect low grade mosaicism and genomic copy number changes below the resolution stated. Balanced rearrangements and Robertsonian translocations will not be detected. This test does not exclude single gene disorders caused by sequence mutations or trinucleotide repeat expansions (such as fragile X syndrome, Huntington disease, some spinocerebellar ataxias, Friedreich ataxia and myotonic dystrophy). Testing for fragile X syndrome should be considered in individuals with developmental delay/ intellectual disability. Further genomic based testing may be considered if there remains a high suspicion of a monogenic disorder. Copy number variants that do not contain genes, are well established polymorphisms, or are assessed as being of unlikely clinical significance (based on “ACMG Technical Standards for the Interpretation and Reporting of Constitutional Copy Number Variants”), will not be reported. The classification is based on the current scientific evidence available at the time of reporting. Reporting of regions of homozygosity (>5Mb) is dependent on referral setting and clinical indication. CNVs that contain autosomal recessive genes will not be reported unless there is specific clinical relevance, high carrier population frequency or a history of consanguinity. Please contact the laboratory if there is a family history of a known recessive disorder or a clinically suspected recessive condition. CNVs involving moderate to high risk cancer susceptibility genes may be reported. However, incidentally ascertained CNVs involving genes assessed as having low to moderate cancer risk will not generally be reported unless they are part of a larger CNV. This testing was performed on a standard SNP microarray platform which may not have sufficient probe coverage to detect clinically relevant CNVs related to this patient’s specific clinical features. Please contact the laboratory if a higher resolution microarray may be required for a specific gene or genetic condition. This test does not exclude the possibility of tissue limited mosaicism and further testing of an alternative tissue may be considered if clinically indicated. Interpretation is based on the UCSC GRCh38/hg38 human reference sequence.*

Validated: 05-Jun-2023 by David Francis

Enquiries: +61 1300 11 8247

END OF TEST REPORT

**FINAL REPORT**

PATHOLOGY REPORT

VCGS
leaders in genetic health

Victorian Clinical Genetics Services
Murdoch Children's Research Institute
Flemington Road, Parkville, VIC 3052 Australia
P +61 1300 118247 F +61 3 8341 6366
W vcgs.org.au ABN 51 007 032 760

LIMS Report #: 1018770

**Patient:** BD009.2 IPSC P11 Bd009.2 Ipsc P11

**DOB:** Unknown
**Sex:** Unknown
**VCGS sample ID:** 23C106576
**Date collected:** 06-Jun-2023
**Date received:** 06-Jun-2023
**Date reported:** 29-Jun-2023
**Source:** Cell Pellet
**Ext. sample ID:**
**Ext. patient ID:**

Dr. Katerina Vlahos
Murdoch Children's Research Institute
Level 5/Flemington Road
PARKVILLE VIC 3052

PATHOLOGY REPORT

### Cytogenetics Laboratory

| | |
|---|---|
| **Clinical details** | Cell line |
| **Specimen source** | Cell Pellet |

#### Molecular karyotype

| | |
|---|---|
| **Array type** | Illumina Infinium GSA-24 v3.0 |
| **Resolution** | 0.50Mb |
| **Reference genome** | GRCh38 / hg38 (Dec 2013) |
| **Molecular karyotype** | arr(X,Y)x1,(1-22)x2 |
| **Result** | NO ANEUPLOIDIES DETECTED |

**Interpretation**

Male molecular karyotype. No aneuploidies were detected in this sample.

*Molecular karyotyping is limited in its ability to detect low grade mosaicism and genomic copy number changes below the resolution stated. Balanced rearrangements and Robertsonian translocations will not be detected. This test does not exclude single gene disorders caused by sequence mutations or trinucleotide repeat expansions (such as fragile X syndrome, Huntington disease, some spinocerebellar ataxias, Friedreich ataxia and myotonic dystrophy). Testing for fragile X syndrome should be considered in individuals with developmental delay/ intellectual disability. Further genomic based testing may be considered if there remains a high suspicion of a monogenic disorder. Copy number variants that do not contain genes, are well established polymorphisms, or are assessed as being of unlikely clinical significance (based on "ACMG Technical Standards for the Interpretation and Reporting of Constitutional Copy Number Variants"), will not be reported. The classification is based on the current scientific evidence available at the time of reporting. Reporting of regions of homozygosity (>5Mb) is dependent on referral setting and clinical indication. CNVs that contain autosomal recessive genes will not be reported unless there is specific clinical relevance, high carrier population frequency or a history of consanguinity. Please contact the laboratory if there is a family history of a known recessive disorder or a clinically suspected recessive condition. CNVs involving moderate to high risk cancer susceptibility genes may be reported. However, incidentally ascertained CNVs involving genes assessed as having low to moderate cancer risk will not generally be reported unless they are part of a larger CNV. This testing was performed on a standard SNP microarray platform which may not have sufficient probe coverage to detect clinically relevant CNVs related to this patient's specific clinical features. Please contact the laboratory if a higher resolution microarray may be required for a specific gene or genetic condition. This test does not exclude the possibility of tissue limited mosaicism and further testing of an alternative tissue may be considered if clinically indicated. Interpretation is based on the UCSC GRCh38/hg38 human reference sequence.*

Validated: 29-Jun-2023 by David Francis

Enquiries: +61 1300 11 8247

**END OF TEST REPORT**

**FINAL REPORT**

Victorian Clinical Genetics Services
Murdoch Children's Research Institute
Flemington Road, Parkville, VIC 3052 Australia
P +61 1300 118247 F +61 3 8341 6366
W vcgs.org.au ABN 51 007 032 760

leaders in genetic health

LIMS Report #: 850338

**Patient:** BD001 P11 Bd001 P11

Dr. Katerina Vlahos
Murdoch Childrens Research Institute
Level 5/Flemington Road
PARKVILLE VIC 3052

**DOB:** Unknown
**Sex:** Unknown
**VCGS sample ID:** 21C111367
**Date collected:** 14-Dec-2021
**Date received:** 14-Dec-2021
**Date reported:** 05-Jan-2022
**Source:** Cell Pellet
**Ext. sample ID:**
**Ext. patient ID:**

### Cytogenetics Laboratory

| | |
|---|---|
| **Clinical details** | Cell line |
| **Specimen source** | Cell Pellet |

#### Molecular karyotype

| | |
|---|---|
| **Array type** | Illumina Infinium GSA-24 v3.0 |
| **Resolution** | 0.50Mb |
| **Assembly** | hg19 / GRCh37 (Feb 2009) |
| **Molecular karyotype** | arr(X,1-22)x2 |
| **Result** | NO ANEUPLOIDIES DETECTED |

**Interpretation**

Female molecular karyotype. No aneuploidies were detected in this sample.

*Molecular karyotyping is limited in its ability to detect low grade mosaicism and genomic copy number changes below the resolution stated. Balanced rearrangements and Robertsonian translocations will not be detected. This test does not exclude single gene disorders caused by sequence mutations or trinucleotide repeat expansions (such as fragile X syndrome, Huntington disease, some spinocerebellar ataxias, Friedreich ataxia and myotonic dystrophy). Testing for fragile X syndrome should be considered in individuals with developmental delay/ intellectual disability. Further genomic based testing may be considered if there remains a high suspicion of a monogenic disorder. Copy number variants that do not contain genes, are well established polymorphisms, or are assessed as being of unlikely clinical significance (based on "ACMG Technical Standards for the Interpretation and Reporting of Constitutional Copy Number Variants"), will not be reported. The classification is based on the current scientific evidence available at the time of reporting. Reporting of regions of homozygosity (>5Mb) is dependent on referral setting and clinical indication. CNVs that contain autosomal recessive genes will not be reported unless there is specific clinical relevance, high carrier population frequency or a history of consanguinity. Please contact the laboratory if there is a family history of a known recessive disorder or a clinically suspected recessive condition. This testing was performed on a standard SNP microarray platform which may not have sufficient probe coverage to detect clinically relevant CNVs related to this patient's specific clinical features. Please contact the laboratory if a higher resolution microarray may be required for a specific gene or genetic condition. This test does not exclude the possibility of tissue limited mosaicism and further testing of an alternative tissue may be considered if clinically indicated. Interpretation is based on the UCSC GRCh37/hg19 human reference sequence.*

Validated: 05-Jan-2022 by David Francis

Enquiries: +61 1300 11 8247

PATHOLOGY REPORT

**END OF TEST REPORT**

**FINAL REPORT**

Victorian Clinical Genetics Services
Murdoch Children's Research Institute
Flemington Road, Parkville, VIC 3052 Australia
P +61 1300 118247 F +61 3 8341 6366
W vcgs.org.au ABN 51 007 032 760

leaders in genetic health

LIMS Report #: 922190

**Patient:** BD004.9 IPSC P10 Bd004.9 Ipsc P10

**DOB:** Unknown
**Sex:** Unknown
**VCGS sample ID:** 22C107158
**Date collected:** 10-Aug-2022
**Date received:** 10-Aug-2022
**Date reported:** 23-Aug-2022
**Source:** Cell Pellet
**Ext. sample ID:**
**Ext. patient ID:**

Dr. Katerina Vlahos
Murdoch Children's Research Institute
Level 5/Flemington Road
PARKVILLE VIC 3052

**CC:** Ips.gene

### Cytogenetics Laboratory

**Clinical details** Cell line
**Specimen source** Cell Pellet

#### Molecular karyotype

| | |
|---|---|
| **Array type** | Illumina Infinium GSA-24 v3.0 |
| **Resolution** | 0.50Mb |
| **Assembly** | hg19 / GRCh37 (Feb 2009) |
| **Molecular karyotype** | arr(X,1-22)x2 |
| **Result** | NO ANEUPLOIDIES DETECTED |

**Interpretation**

Female molecular karyotype. No aneuploidies were detected in this sample.

*Molecular karyotyping is limited in its ability to detect low grade mosaicism and genomic copy number changes below the resolution stated. Balanced rearrangements and Robertsonian translocations will not be detected. This test does not exclude single gene disorders caused by sequence mutations or trinucleotide repeat expansions (such as fragile X syndrome, Huntington disease, some spinocerebellar ataxias, Friedreich ataxia and myotonic dystrophy). Testing for fragile X syndrome should be considered in individuals with developmental delay/ intellectual disability. Further genomic based testing may be considered if there remains a high suspicion of a monogenic disorder. Copy number variants that do not contain genes, are well established polymorphisms, or are assessed as being of unlikely clinical significance (based on “ACMG Technical Standards for the Interpretation and Reporting of Constitutional Copy Number Variants”), will not be reported. The classification is based on the current scientific evidence available at the time of reporting. Reporting of regions of homozygosity (>5Mb) is dependent on referral setting and clinical indication. CNVs that contain autosomal recessive genes will not be reported unless there is specific clinical relevance, high carrier population frequency or a history of consanguinity. Please contact the laboratory if there is a family history of a known recessive disorder or a clinically suspected recessive condition. This testing was performed on a standard SNP microarray platform which may not have sufficient probe coverage to detect clinically relevant CNVs related to this patient’s specific clinical features. Please contact the laboratory if a higher resolution microarray may be required for a specific gene or genetic condition. This test does not exclude the possibility of tissue limited mosaicism and further testing of an alternative tissue may be considered if clinically indicated. Interpretation is based on the UCSC GRCh37/hg19 human reference sequence.*

Validated: 23-Aug-2022 by David Francis

Enquiries: +61 1300 11 8247

PATHOLOGY REPORT

**END OF TEST REPORT**

**FINAL REPORT**

Victorian Clinical Genetics Services
Murdoch Children's Research Institute
Flemington Road, Parkville, VIC 3052 Australia
P +61 1300 118247 F +61 3 8341 6366
W vcgs.org.au ABN 51 007 032 760

leaders in genetic health

LIMS Report #: 850376

**Patient:** BD012.2 P11 Bd012.2 P11

**DOB:** Unknown
**Sex:** Unknown
**VCGS sample ID:** 21C111373
**Date collected:** 14-Dec-2021
**Date received:** 14-Dec-2021
**Date reported:** 05-Jan-2022
**Source:** Cell Pellet
**Ext. sample ID:**
**Ext. patient ID:**

Dr. Katerina Vlahos
Murdoch Childrens Research Institute
Level 5/Flemington Road
PARKVILLE VIC 3052

### Cytogenetics Laboratory

| | |
|---|---|
| **Clinical details** | Cell line |
| **Specimen source** | Cell Pellet |

#### Molecular karyotype

| | |
|---|---|
| **Array type** | Illumina Infinium GSA-24 v3.0 |
| **Resolution** | 0.50Mb |
| **Assembly** | hg19 / GRCh37 (Feb 2009) |
| **Molecular karyotype** | arr(X,Y)x1,(1-22)x2 |
| **Result** | NO ANEUPLOIDIES DETECTED |

**Interpretation**

Male molecular karyotype. No aneuploidies were detected in this sample.

*Molecular karyotyping is limited in its ability to detect low grade mosaicism and genomic copy number changes below the resolution stated. Balanced rearrangements and Robertsonian translocations will not be detected. This test does not exclude single gene disorders caused by sequence mutations or trinucleotide repeat expansions (such as fragile X syndrome, Huntington disease, some spinocerebellar ataxias, Friedreich ataxia and myotonic dystrophy). Testing for fragile X syndrome should be considered in individuals with developmental delay/ intellectual disability. Further genomic based testing may be considered if there remains a high suspicion of a monogenic disorder. Copy number variants that do not contain genes, are well established polymorphisms, or are assessed as being of unlikely clinical significance (based on "ACMG Technical Standards for the Interpretation and Reporting of Constitutional Copy Number Variants"), will not be reported. The classification is based on the current scientific evidence available at the time of reporting. Reporting of regions of homozygosity (>5Mb) is dependent on referral setting and clinical indication. CNVs that contain autosomal recessive genes will not be reported unless there is specific clinical relevance, high carrier population frequency or a history of consanguinity. Please contact the laboratory if there is a family history of a known recessive disorder or a clinically suspected recessive condition. This testing was performed on a standard SNP microarray platform which may not have sufficient probe coverage to detect clinically relevant CNVs related to this patient's specific clinical features. Please contact the laboratory if a higher resolution microarray may be required for a specific gene or genetic condition. This test does not exclude the possibility of tissue limited mosaicism and further testing of an alternative tissue may be considered if clinically indicated. Interpretation is based on the UCSC GRCh37/hg19 human reference sequence.*

Validated: 05-Jan-2022 by David Francis

Enquiries: +61 1300 11 8247

PATHOLOGY REPORT

**END OF TEST REPORT**

**FINAL REPORT**

Victorian Clinical Genetics Services
Murdoch Children's Research Institute
Flemington Road, Parkville, VIC 3052 Australia
P +61 1300 118247 F +61 3 8341 6366
W vcgs.org.au ABN 51 007 032 760

VCGS leaders in genetic health

LIMS Report #: 1018767

**Patient:** BD005.5 IPSC P11 Bd005.5 Ipsc P11

**DOB:** Unknown
**Sex:** Unknown
**VCGS sample ID:** 23C106574
**Date collected:** 06-Jun-2023
**Date received:** 06-Jun-2023
**Date reported:** 29-Jun-2023
**Source:** Cell Pellet
**Ext. sample ID:**
**Ext. patient ID:**

Dr. Katerina Vlahos
Murdoch Children's Research Institute
Level 5/Flemington Road
PARKVILLE VIC 3052

**CC:** Ips.gene

### Cytogenetics Laboratory

| | |
|---|---|
| **Clinical details** | Cell line |
| **Specimen source** | Cell Pellet |

#### Molecular karyotype

| | |
|---|---|
| **Array type** | Illumina Infinium GSA-24 v3.0 |
| **Resolution** | 0.50Mb |
| **Reference genome** | GRCh38 / hg38 (Dec 2013) |
| **Molecular karyotype** | arr(X,Y)x1,(1-22)x2 |
| **Result** | NO ANEUPLOIDIES DETECTED |

**Interpretation**

Male molecular karyotype. No aneuploidies were detected in this sample.

*Molecular karyotyping is limited in its ability to detect low grade mosaicism and genomic copy number changes below the resolution stated. Balanced rearrangements and Robertsonian translocations will not be detected. This test does not exclude single gene disorders caused by sequence mutations or trinucleotide repeat expansions (such as fragile X syndrome, Huntington disease, some spinocerebellar ataxias, Friedreich ataxia and myotonic dystrophy). Testing for fragile X syndrome should be considered in individuals with developmental delay/ intellectual disability. Further genomic based testing may be considered if there remains a high suspicion of a monogenic disorder. Copy number variants that do not contain genes, are well established polymorphisms, or are assessed as being of unlikely clinical significance (based on "ACMG Technical Standards for the Interpretation and Reporting of Constitutional Copy Number Variants"), will not be reported. The classification is based on the current scientific evidence available at the time of reporting. Reporting of regions of homozygosity (>5Mb) is dependent on referral setting and clinical indication. CNVs that contain autosomal recessive genes will not be reported unless there is specific clinical relevance, high carrier population frequency or a history of consanguinity. Please contact the laboratory if there is a family history of a known recessive disorder or a clinically suspected recessive condition. CNVs involving moderate to high risk cancer susceptibility genes may be reported. However, incidentally ascertained CNVs involving genes assessed as having low to moderate cancer risk will not generally be reported unless they are part of a larger CNV. This testing was performed on a standard SNP microarray platform which may not have sufficient probe coverage to detect clinically relevant CNVs related to this patient's specific clinical features. Please contact the laboratory if a higher resolution microarray may be required for a specific gene or genetic condition. This test does not exclude the possibility of tissue limited mosaicism and further testing of an alternative tissue may be considered if clinically indicated. Interpretation is based on the UCSC GRCh38/hg38 human reference sequence.*

Validated: 29-Jun-2023 by David Francis

Enquiries: +61 1300 11 8247

**END OF TEST REPORT**

**FINAL REPORT**

PATHOLOGY REPORT

Victorian Clinical Genetics Services
Murdoch Children's Research Institute
Flemington Road, Parkville, VIC 3052 Australia
P +61 1300 118247 F +61 3 8341 6366
W vcgs.org.au ABN 51 007 032 760

leaders in genetic health

LIMS Report #: 901661

**Patient:** BD002.3 P10 Bd002.3 P10

Dr. Katerina Vlahos
Murdoch Children's Research Institute
Level 5/Flemington Road
PARKVILLE VIC 3052

**DOB:** Unknown
**Sex:** Unknown
**VCGS sample ID:** 22C103779
**Date collected:** 05-May-2022
**Date received:** 05-May-2022
**Date reported:** 20-Jun-2022
**Source:** Cell Pellet
**Ext. sample ID:**
**Ext. patient ID:**

**CC:** Ips.gene

### Cytogenetics Laboratory

| | |
|---|---|
| **Clinical details** | Cell line |
| **Specimen source** | Cell Pellet |

#### Molecular karyotype

| | |
|---|---|
| **Array type** | Illumina Infinium GSA-24 v3.0 |
| **Resolution** | 0.50Mb |
| **Assembly** | hg19 / GRCh37 (Feb 2009) |
| **Molecular karyotype** | arr(X,1-22)x2 |
| **Result** | NO ANEUPLOIDIES DETECTED |

**Interpretation**

Female molecular karyotype. No aneuploidies were detected in this sample.

*Molecular karyotyping is limited in its ability to detect low grade mosaicism and genomic copy number changes below the resolution stated. Balanced rearrangements and Robertsonian translocations will not be detected. This test does not exclude single gene disorders caused by sequence mutations or trinucleotide repeat expansions (such as fragile X syndrome, Huntington disease, some spinocerebellar ataxias, Friedreich ataxia and myotonic dystrophy). Testing for fragile X syndrome should be considered in individuals with developmental delay/ intellectual disability. Further genomic based testing may be considered if there remains a high suspicion of a monogenic disorder. Copy number variants that do not contain genes, are well established polymorphisms, or are assessed as being of unlikely clinical significance (based on "ACMG Technical Standards for the Interpretation and Reporting of Constitutional Copy Number Variants"), will not be reported. The classification is based on the current scientific evidence available at the time of reporting. Reporting of regions of homozygosity (>5Mb) is dependent on referral setting and clinical indication. CNVs that contain autosomal recessive genes will not be reported unless there is specific clinical relevance, high carrier population frequency or a history of consanguinity. Please contact the laboratory if there is a family history of a known recessive disorder or a clinically suspected recessive condition. This testing was performed on a standard SNP microarray platform which may not have sufficient probe coverage to detect clinically relevant CNVs related to this patient's specific clinical features. Please contact the laboratory if a higher resolution microarray may be required for a specific gene or genetic condition. This test does not exclude the possibility of tissue limited mosaicism and further testing of an alternative tissue may be considered if clinically indicated. Interpretation is based on the UCSC GRCh37/hg19 human reference sequence.*

Validated: 20-Jun-2022 by David Francis

Enquiries: +61 1300 11 8247

**END OF TEST REPORT**

**FINAL REPORT**

PATHOLOGY REPORT

Victorian Clinical Genetics Services
Murdoch Children's Research Institute
Flemington Road, Parkville, VIC 3052 Australia
P +61 1300 118247 F +61 3 8341 6366
W vcgs.org.au ABN 51 007 032 760

LIMS Report #: 850370

**Patient:** BD010.5 P10 Bd010.5 P10

Dr. Katerina Vlahos
Murdoch Childrens Research Institute
Level 5/Flemington Road
PARKVILLE VIC 3052

**DOB:** Unknown
**Sex:** Unknown
**VCGS sample ID:** 21C111372
**Date collected:** 14-Dec-2021
**Date received:** 14-Dec-2021
**Date reported:** 05-Jan-2022
**Source:** Cell Pellet
**Ext. sample ID:**
**Ext. patient ID:**

### Cytogenetics Laboratory

| | |
|---|---|
| **Clinical details** | Cell line |
| **Specimen source** | Cell Pellet |

#### Molecular karyotype

| | |
|---|---|
| **Array type** | Illumina Infinium GSA-24 v3.0 |
| **Resolution** | 0.50Mb |
| **Assembly** | hg19 / GRCh37 (Feb 2009) |
| **Molecular karyotype** | arr(X,1-22)x2 |
| **Result** | NO ANEUPLOIDIES DETECTED |

**Interpretation**
Female molecular karyotype. No aneuploidies were detected in this sample.

*Molecular karyotyping is limited in its ability to detect low grade mosaicism and genomic copy number changes below the resolution stated. Balanced rearrangements and Robertsonian translocations will not be detected. This test does not exclude single gene disorders caused by sequence mutations or trinucleotide repeat expansions (such as fragile X syndrome, Huntington disease, some spinocerebellar ataxias, Friedreich ataxia and myotonic dystrophy). Testing for fragile X syndrome should be considered in individuals with developmental delay/ intellectual disability. Further genomic based testing may be considered if there remains a high suspicion of a monogenic disorder. Copy number variants that do not contain genes, are well established polymorphisms, or are assessed as being of unlikely clinical significance (based on "ACMG Technical Standards for the Interpretation and Reporting of Constitutional Copy Number Variants"), will not be reported. The classification is based on the current scientific evidence available at the time of reporting. Reporting of regions of homozygosity (>5Mb) is dependent on referral setting and clinical indication. CNVs that contain autosomal recessive genes will not be reported unless there is specific clinical relevance, high carrier population frequency or a history of consanguinity. Please contact the laboratory if there is a family history of a known recessive disorder or a clinically suspected recessive condition. This testing was performed on a standard SNP microarray platform which may not have sufficient probe coverage to detect clinically relevant CNVs related to this patient's specific clinical features. Please contact the laboratory if a higher resolution microarray may be required for a specific gene or genetic condition. This test does not exclude the possibility of tissue limited mosaicism and further testing of an alternative tissue may be considered if clinically indicated. Interpretation is based on the UCSC GRCh37/hg19 human reference sequence.*

Validated: 05-Jan-2022 by David Francis

Enquiries: +61 1300 11 8247

PATHOLOGY REPORT

**END OF TEST REPORT**

**FINAL REPORT**

Victorian Clinical Genetics Services
Murdoch Children's Research Institute
Flemington Road, Parkville, VIC 3052 Australia
P +61 1300 118247 F +61 3 8341 6366
W vcgs.org.au ABN 51 007 032 760

leaders in genetic health

LIMS Report #: 850369

**Patient:** BD008.2 P10 Bd008.2 P10

**DOB:** Unknown
**Sex:** Unknown
**VCGS sample ID:** 21C111371
**Date collected:** 14-Dec-2021
**Date received:** 14-Dec-2021
**Date reported:** 05-Jan-2022
**Source:** Cell Pellet
**Ext. sample ID:**
**Ext. patient ID:**

Dr. Katerina Vlahos
Murdoch Childrens Research Institute
Level 5/Flemington Road
PARKVILLE VIC 3052

### Cytogenetics Laboratory

| | |
|---|---|
| **Clinical details** | Cell line |
| **Specimen source** | Cell Pellet |

#### Molecular karyotype

| | |
|---|---|
| **Array type** | Illumina Infinium GSA-24 v3.0 |
| **Resolution** | 0.50Mb |
| **Assembly** | hg19 / GRCh37 (Feb 2009) |
| **Molecular karyotype** | arr(X,1-22)x2 |
| **Result** | NO ANEUPLOIDIES DETECTED |

**Interpretation**

Female molecular karyotype. No aneuploidies were detected in this sample.

*Molecular karyotyping is limited in its ability to detect low grade mosaicism and genomic copy number changes below the resolution stated. Balanced rearrangements and Robertsonian translocations will not be detected. This test does not exclude single gene disorders caused by sequence mutations or trinucleotide repeat expansions (such as fragile X syndrome, Huntington disease, some spinocerebellar ataxias, Friedreich ataxia and myotonic dystrophy). Testing for fragile X syndrome should be considered in individuals with developmental delay/ intellectual disability. Further genomic based testing may be considered if there remains a high suspicion of a monogenic disorder. Copy number variants that do not contain genes, are well established polymorphisms, or are assessed as being of unlikely clinical significance (based on "ACMG Technical Standards for the Interpretation and Reporting of Constitutional Copy Number Variants"), will not be reported. The classification is based on the current scientific evidence available at the time of reporting. Reporting of regions of homozygosity (>5Mb) is dependent on referral setting and clinical indication. CNVs that contain autosomal recessive genes will not be reported unless there is specific clinical relevance, high carrier population frequency or a history of consanguinity. Please contact the laboratory if there is a family history of a known recessive disorder or a clinically suspected recessive condition. This testing was performed on a standard SNP microarray platform which may not have sufficient probe coverage to detect clinically relevant CNVs related to this patient's specific clinical features. Please contact the laboratory if a higher resolution microarray may be required for a specific gene or genetic condition. This test does not exclude the possibility of tissue limited mosaicism and further testing of an alternative tissue may be considered if clinically indicated. Interpretation is based on the UCSC GRCh37/hg19 human reference sequence.*

Validated: 05-Jan-2022 by David Francis

Enquiries: +61 1300 11 8247

PATHOLOGY REPORT

**END OF TEST REPORT**

**FINAL REPORT**

Victorian Clinical Genetics Services
Murdoch Children's Research Institute
Flemington Road, Parkville, VIC 3052 Australia
P +61 1300 118247 F +61 3 8341 6366
W vcgs.org.au ABN 51 007 032 760

LIMS Report #: 901645

**Patient:** BD003.1 P12 Bd003.1 P12

**DOB:** Unknown
**Sex:** Unknown
**VCGS sample ID:** 22C103775
**Date collected:** 05-May-2022
**Date received:** 05-May-2022
**Date reported:** 20-Jun-2022
**Source:** Cell Pellet
**Ext. sample ID:**
**Ext. patient ID:**

Dr. Katerina Vlahos
Murdoch Children's Research Institute
Level 5/Flemington Road
PARKVILLE VIC 3052

**CC:** Ips.gene

### Cytogenetics Laboratory

| | |
|---|---|
| **Clinical details** | Cell line |
| **Specimen source** | Cell Pellet |

#### Molecular karyotype

| | |
|---|---|
| **Array type** | Illumina Infinium GSA-24 v3.0 |
| **Resolution** | 0.50Mb |
| **Assembly** | hg19 / GRCh37 (Feb 2009) |
| **Molecular karyotype** | arr(X,1-22)x2 |
| **Result** | NO ANEUPLOIDIES DETECTED |

**Interpretation**

Female molecular karyotype. No aneuploidies were detected in this sample.

*Molecular karyotyping is limited in its ability to detect low grade mosaicism and genomic copy number changes below the resolution stated. Balanced rearrangements and Robertsonian translocations will not be detected. This test does not exclude single gene disorders caused by sequence mutations or trinucleotide repeat expansions (such as fragile X syndrome, Huntington disease, some spinocerebellar ataxias, Friedreich ataxia and myotonic dystrophy). Testing for fragile X syndrome should be considered in individuals with developmental delay/ intellectual disability. Further genomic based testing may be considered if there remains a high suspicion of a monogenic disorder. Copy number variants that do not contain genes, are well established polymorphisms, or are assessed as being of unlikely clinical significance (based on “ACMG Technical Standards for the Interpretation and Reporting of Constitutional Copy Number Variants”), will not be reported. The classification is based on the current scientific evidence available at the time of reporting. Reporting of regions of homozygosity (>5Mb) is dependent on referral setting and clinical indication. CNVs that contain autosomal recessive genes will not be reported unless there is specific clinical relevance, high carrier population frequency or a history of consanguinity. Please contact the laboratory if there is a family history of a known recessive disorder or a clinically suspected recessive condition. This testing was performed on a standard SNP microarray platform which may not have sufficient probe coverage to detect clinically relevant CNVs related to this patient’s specific clinical features. Please contact the laboratory if a higher resolution microarray may be required for a specific gene or genetic condition. This test does not exclude the possibility of tissue limited mosaicism and further testing of an alternative tissue may be considered if clinically indicated. Interpretation is based on the UCSC GRCh37/hg19 human reference sequence.*

Validated: 20-Jun-2022 by David Francis

Enquiries: +61 1300 11 8247

PATHOLOGY REPORT

**END OF TEST REPORT**

**FINAL REPORT**

Victorian Clinical Genetics Services
Murdoch Children's Research Institute
Flemington Road, Parkville, VIC 3052 Australia
P +61 1300 118247 F +61 3 8341 6366
W vcgs.org.au ABN 51 007 032 760

leaders in genetic health

LIMS Report #: 850368

**Patient:** BD007.8 P11 Bd007.8 P11

**DOB:** Unknown
**Sex:** Unknown
**VCGS sample ID:** 21C111370
**Date collected:** 14-Dec-2021
**Date received:** 14-Dec-2021
**Date reported:** 05-Jan-2022
**Source:** Cell Pellet
**Ext. sample ID:**
**Ext. patient ID:**

Dr. Katerina Vlahos
Murdoch Childrens Research Institute
Level 5/Flemington Road
PARKVILLE VIC 3052

### Cytogenetics Laboratory

| | |
|---|---|
| **Clinical details** | Cell line |
| **Specimen source** | Cell Pellet |

#### Molecular karyotype

| | |
|---|---|
| **Array type** | Illumina Infinium GSA-24 v3.0 |
| **Resolution** | 0.50Mb |
| **Assembly** | hg19 / GRCh37 (Feb 2009) |
| **Molecular karyotype** | arr(X,1-22)x2 |
| **Result** | NO ANEUPLOIDIES DETECTED |

**Interpretation**

Female molecular karyotype. No aneuploidies were detected in this sample.

*Molecular karyotyping is limited in its ability to detect low grade mosaicism and genomic copy number changes below the resolution stated. Balanced rearrangements and Robertsonian translocations will not be detected. This test does not exclude single gene disorders caused by sequence mutations or trinucleotide repeat expansions (such as fragile X syndrome, Huntington disease, some spinocerebellar ataxias, Friedreich ataxia and myotonic dystrophy). Testing for fragile X syndrome should be considered in individuals with developmental delay/ intellectual disability. Further genomic based testing may be considered if there remains a high suspicion of a monogenic disorder. Copy number variants that do not contain genes, are well established polymorphisms, or are assessed as being of unlikely clinical significance (based on "ACMG Technical Standards for the Interpretation and Reporting of Constitutional Copy Number Variants"), will not be reported. The classification is based on the current scientific evidence available at the time of reporting. Reporting of regions of homozygosity (>5Mb) is dependent on referral setting and clinical indication. CNVs that contain autosomal recessive genes will not be reported unless there is specific clinical relevance, high carrier population frequency or a history of consanguinity. Please contact the laboratory if there is a family history of a known recessive disorder or a clinically suspected recessive condition. This testing was performed on a standard SNP microarray platform which may not have sufficient probe coverage to detect clinically relevant CNVs related to this patient's specific clinical features. Please contact the laboratory if a higher resolution microarray may be required for a specific gene or genetic condition. This test does not exclude the possibility of tissue limited mosaicism and further testing of an alternative tissue may be considered if clinically indicated. Interpretation is based on the UCSC GRCh37/hg19 human reference sequence.*

Validated: 05-Jan-2022 by David Francis

Enquiries: +61 1300 11 8247

PATHOLOGY REPORT

**END OF TEST REPORT**

**FINAL REPORT**

Victorian Clinical Genetics Services
Murdoch Children's Research Institute
Flemington Road, Parkville, VIC 3052 Australia
P +61 1300 118247 F +61 3 8341 6366
W vcgs.org.au ABN 51 007 032 760

VCGS leaders in genetic health

LIMS Report #: 1018769

**Patient:** BD017.7 IPSC P11 Bd017.7 Ipsc P11

**DOB:** Unknown
**Sex:** Unknown
**VCGS sample ID:** 23C106575
**Date collected:** 06-Jun-2023
**Date received:** 06-Jun-2023
**Date reported:** 29-Jun-2023
**Source:** Cell Pellet
**Ext. sample ID:**
**Ext. patient ID:**

Dr. Katerina Vlahos
Murdoch Children's Research Institute
Level 5/Flemington Road
PARKVILLE VIC 3052

**CC:** Ips.gene

### Cytogenetics Laboratory

| | |
|---|---|
| **Clinical details** | Cell line |
| **Specimen source** | Cell Pellet |

#### Molecular karyotype

| | |
|---|---|
| **Array type** | Illumina Infinium GSA-24 v3.0 |
| **Resolution** | 0.50Mb |
| **Reference genome** | GRCh38 / hg38 (Dec 2013) |
| **Molecular karyotype** | arr(X,1-22)x2 |
| **Result** | NO ANEUPLOIDIES DETECTED |

**Interpretation**

Female molecular karyotype. No aneuploidies were detected in this sample.

*Molecular karyotyping is limited in its ability to detect low grade mosaicism and genomic copy number changes below the resolution stated. Balanced rearrangements and Robertsonian translocations will not be detected. This test does not exclude single gene disorders caused by sequence mutations or trinucleotide repeat expansions (such as fragile X syndrome, Huntington disease, some spinocerebellar ataxias, Friedreich ataxia and myotonic dystrophy). Testing for fragile X syndrome should be considered in individuals with developmental delay/ intellectual disability. Further genomic based testing may be considered if there remains a high suspicion of a monogenic disorder. Copy number variants that do not contain genes, are well established polymorphisms, or are assessed as being of unlikely clinical significance (based on "ACMG Technical Standards for the Interpretation and Reporting of Constitutional Copy Number Variants"), will not be reported. The classification is based on the current scientific evidence available at the time of reporting. Reporting of regions of homozygosity (>5Mb) is dependent on referral setting and clinical indication. CNVs that contain autosomal recessive genes will not be reported unless there is specific clinical relevance, high carrier population frequency or a history of consanguinity. Please contact the laboratory if there is a family history of a known recessive disorder or a clinically suspected recessive condition. CNVs involving moderate to high risk cancer susceptibility genes may be reported. However, incidentally ascertained CNVs involving genes assessed as having low to moderate cancer risk will not generally be reported unless they are part of a larger CNV. This testing was performed on a standard SNP microarray platform which may not have sufficient probe coverage to detect clinically relevant CNVs related to this patient's specific clinical features. Please contact the laboratory if a higher resolution microarray may be required for a specific gene or genetic condition. This test does not exclude the possibility of tissue limited mosaicism and further testing of an alternative tissue may be considered if clinically indicated. Interpretation is based on the UCSC GRCh38/hg38 human reference sequence.*

Validated: 29-Jun-2023 by David Francis

Enquiries: +61 1300 11 8247

**END OF TEST REPORT**

**FINAL REPORT**

PATHOLOGY REPORT

VCGS
leaders in genetic health

Victorian Clinical Genetics Services
Murdoch Children's Research Institute
Flemington Road, Parkville, VIC 3052 Australia
P +61 1300 118247 F +61 3 8341 6366
W vcgs.org.au ABN 51 007 032 760

LIMS Report #: 1018766

**Patient:** BD013.3 IPSC P11 Bd013.3 Ipsc P11

**DOB:** Unknown
**Sex:** Unknown
**VCGS sample ID:** 23C106573
**Date collected:** 06-Jun-2023
**Date received:** 06-Jun-2023
**Date reported:** 29-Jun-2023
**Source:** Cell Pellet
**Ext. sample ID:**
**Ext. patient ID:**

Dr. Katerina Vlahos
Murdoch Children's Research Institute
Level 5/Flemington Road
PARKVILLE VIC 3052

**CC:** Ips.gene

PATHOLOGY REPORT

### Cytogenetics Laboratory

| | |
|---|---|
| **Clinical details** | Cell line |
| **Specimen source** | Cell Pellet |

#### Molecular karyotype

| | |
|---|---|
| **Array type** | Illumina Infinium GSA-24 v3.0 |
| **Resolution** | 0.50Mb |
| **Reference genome** | GRCh38 / hg38 (Dec 2013) |
| **Molecular karyotype** | arr(X,1-22)x2 |
| **Result** | NO ANEUPLOIDIES DETECTED |

**Interpretation**

Female molecular karyotype. No aneuploidies were detected in this sample.

*Molecular karyotyping is limited in its ability to detect low grade mosaicism and genomic copy number changes below the resolution stated. Balanced rearrangements and Robertsonian translocations will not be detected. This test does not exclude single gene disorders caused by sequence mutations or trinucleotide repeat expansions (such as fragile X syndrome, Huntington disease, some spinocerebellar ataxias, Friedreich ataxia and myotonic dystrophy). Testing for fragile X syndrome should be considered in individuals with developmental delay/ intellectual disability. Further genomic based testing may be considered if there remains a high suspicion of a monogenic disorder. Copy number variants that do not contain genes, are well established polymorphisms, or are assessed as being of unlikely clinical significance (based on "ACMG Technical Standards for the Interpretation and Reporting of Constitutional Copy Number Variants"), will not be reported. The classification is based on the current scientific evidence available at the time of reporting. Reporting of regions of homozygosity (>5Mb) is dependent on referral setting and clinical indication. CNVs that contain autosomal recessive genes will not be reported unless there is specific clinical relevance, high carrier population frequency or a history of consanguinity. Please contact the laboratory if there is a family history of a known recessive disorder or a clinically suspected recessive condition. CNVs involving moderate to high risk cancer susceptibility genes may be reported. However, incidentally ascertained CNVs involving genes assessed as having low to moderate cancer risk will not generally be reported unless they are part of a larger CNV. This testing was performed on a standard SNP microarray platform which may not have sufficient probe coverage to detect clinically relevant CNVs related to this patient's specific clinical features. Please contact the laboratory if a higher resolution microarray may be required for a specific gene or genetic condition. This test does not exclude the possibility of tissue limited mosaicism and further testing of an alternative tissue may be considered if clinically indicated. Interpretation is based on the UCSC GRCh38/hg38 human reference sequence.*

Validated: 29-Jun-2023 by David Francis

Enquiries: +61 1300 11 8247

**END OF TEST REPORT**

**FINAL REPORT**

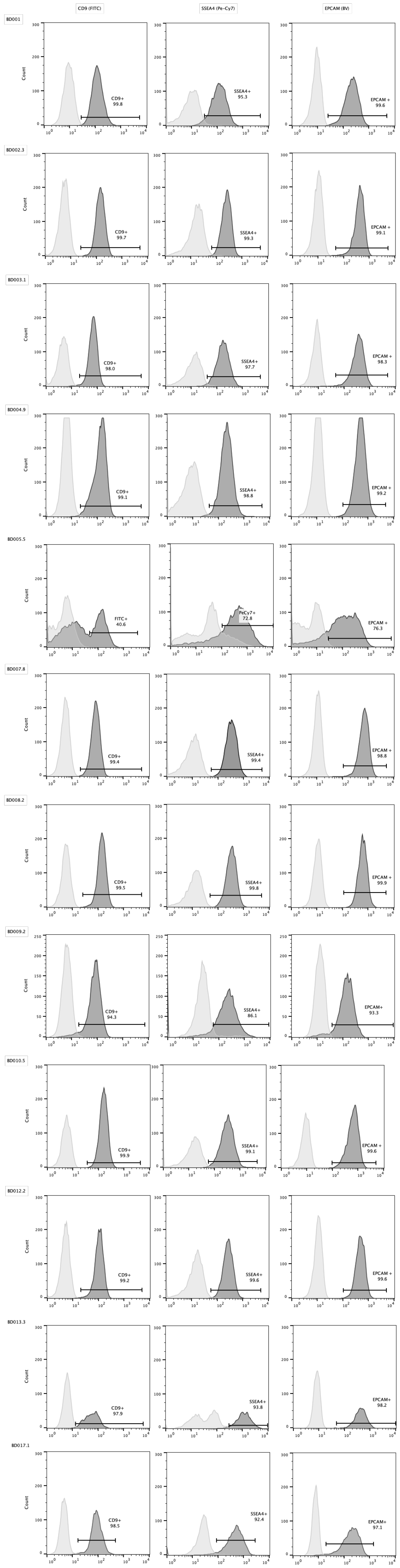

CD9 (FITC)
SSEA4 (Pe-Cy7)
EPCAM (BV)
BD001
CD9+ 99.8
SSEA4+ 95.3
EPCAM + 99.6
BD002.3
CD9+ 99.7
SSEA4+ 99.3
EPCAM + 99.1
BD003.1
CD9+ 98.0
SSEA4+ 97.7
EPCAM + 98.3
BD004.9
CD9+ 99.1
SSEA4+ 98.8
EPCAM + 99.2
BD005.5
FITC+ 40.6
PeCy7+ 72.8
EPCAM + 76.3
BD007.8
CD9+ 99.4
SSEA4+ 99.4
EPCAM + 98.8
BD008.2
CD9+ 99.5
SSEA4+ 99.8
EPCAM + 99.9
BD009.2
CD9+ 94.3
SSEA4+ 86.1
EPCAM+ 93.3
BD010.5
CD9+ 99.9
SSEA4+ 99.1
EPCAM + 99.6
BD012.2
CD9+ 99.2
SSEA4+ 99.6
EPCAM + 99.6
BD013.3
CD9+ 97.9
SSEA4+ 93.8
EPCAM+ 98.2
BD017.1
CD9+ 98.5
SSEA4+ 92.4
EPCAM+ 97.1
Count

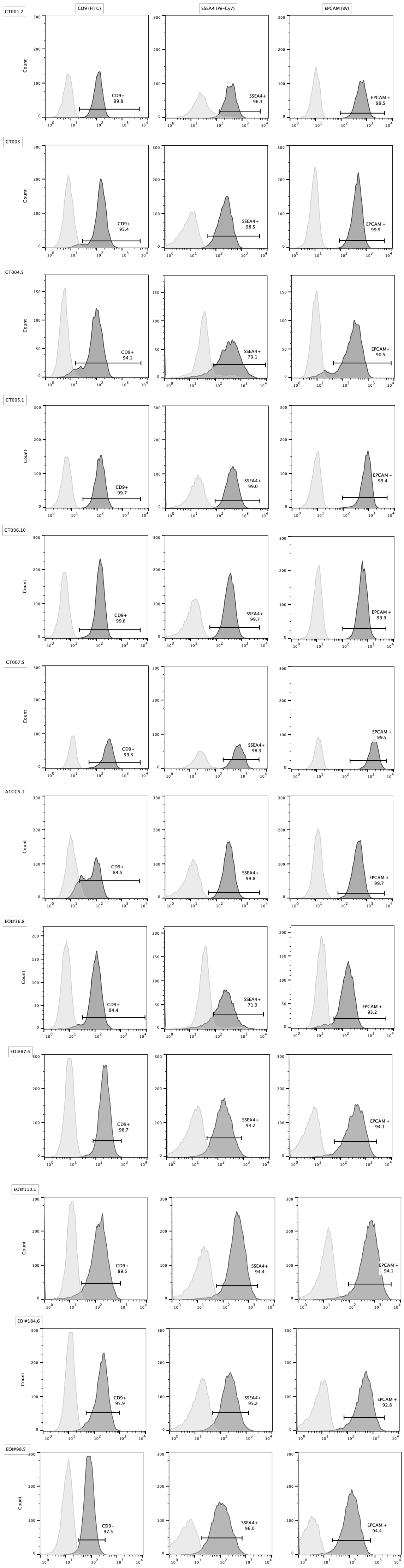

CD9 (FITC)
SSEA4 (Pe-Cy7)
EPCAM (BV)
Count
CT001.7
CD9+ 99.8
SSEA4+ 96.3
EPCAM + 99.5
CT003
CD9+ 95.4
SSEA4+ 98.5
EPCAM + 99.5
CT004.5
CD9+ 94.1
SSEA4+ 79.1
EPCAM+ 90.5
CT005.1
CD9+ 99.7
SSEA4+ 99.0
EPCAM + 99.4
CT006.10
CD9+ 99.6
SSEA4+ 99.7
EPCAM + 99.9
CT007.5
CD9+ 99.3
SSEA4+ 98.3
EPCAM + 99.5
ATCC5.1
CD9+ 84.5
SSEA4+ 99.8
EPCAM + 99.7
EOI#36.8
CD9+ 94.4
SSEA4+ 71.3
EPCAM + 93.2
EOI#87.4
CD9+ 96.7
SSEA4+ 94.2
EPCAM + 94.1
EOI#110.1
CD9+ 89.5
SSEA4+ 94.4
EPCAM + 94.1
EOI#184.6
CD9+ 95.9
SSEA4+ 95.2
EPCAM + 92.8
EOI#98.5
CD9+ 97.5
SSEA4+ 96.0
EPCAM + 94.4

Supplementary figure 1: iPSCs

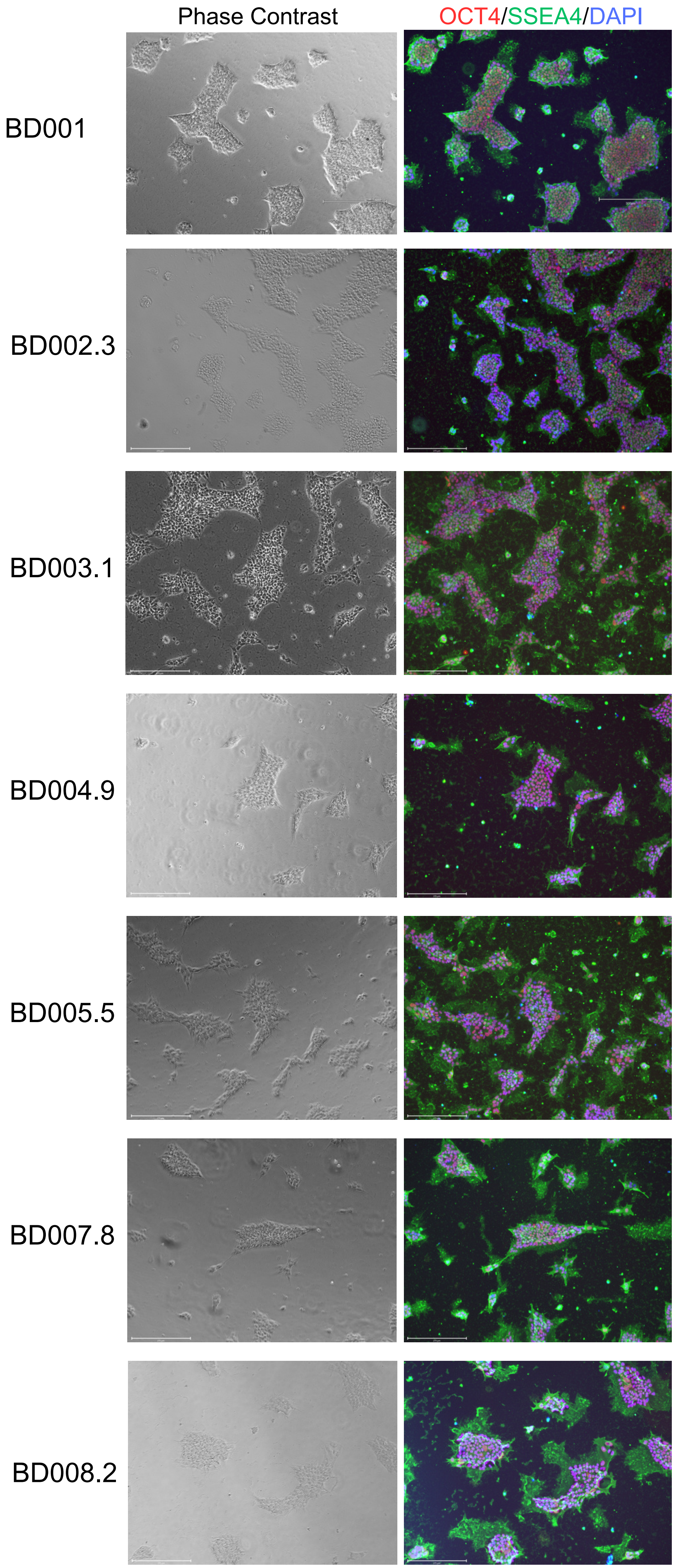

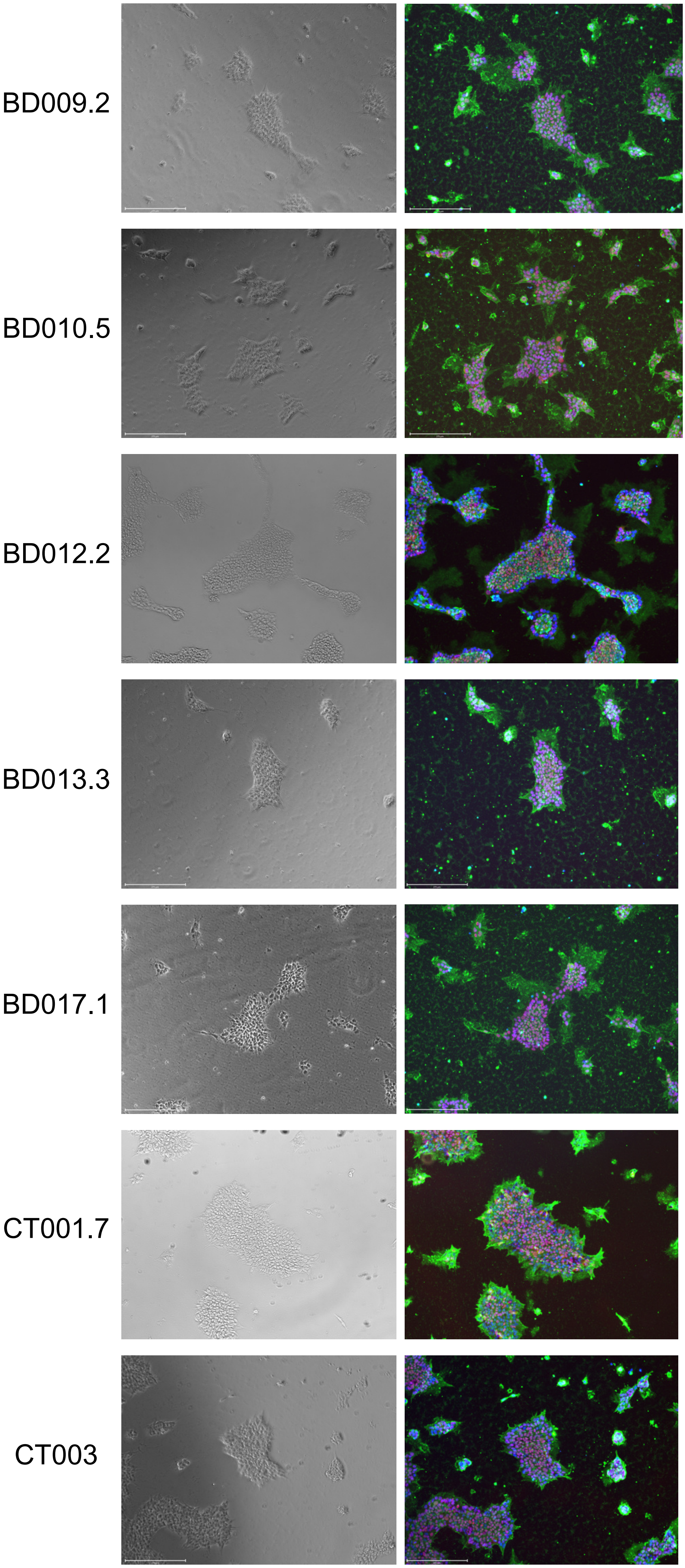

BD009.2
BD010.5
BD012.2
BD013.3
BD017.1
CT001.7
CT003

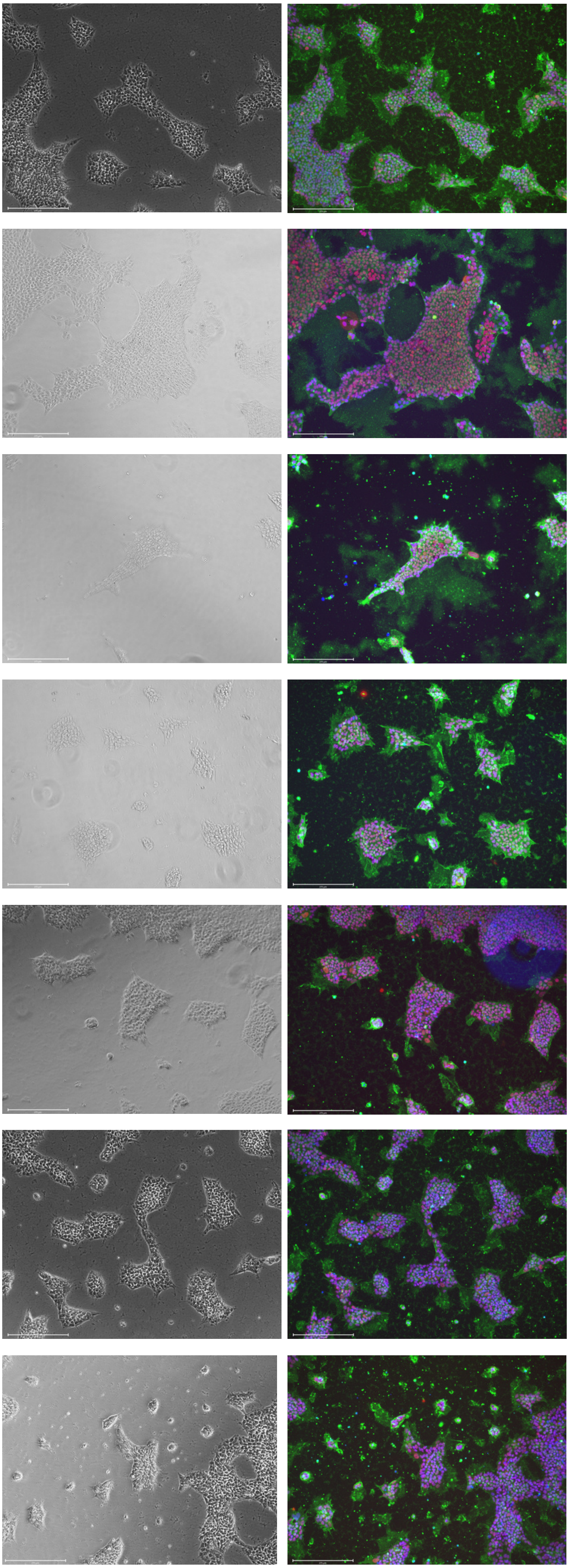

CT004.5

CT005.1

CT006.10

CT007.5

ATCC5.1

EOI#36.8

EOI#98.5

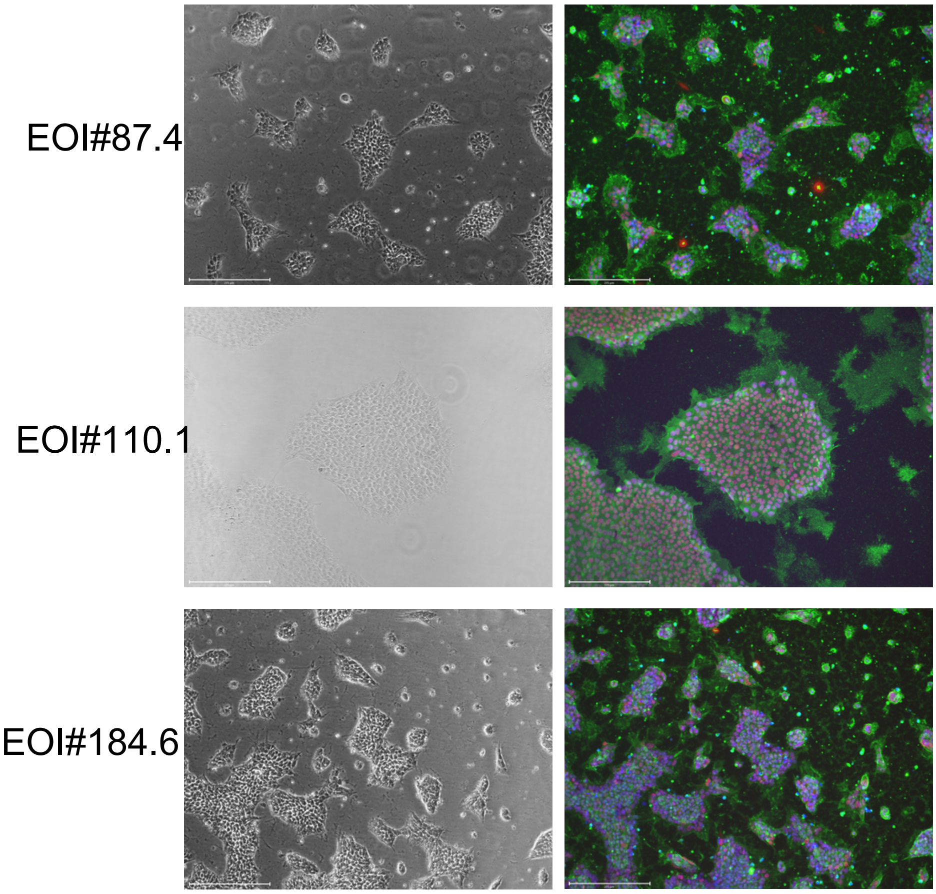


Images of the 24 iPSC lines generated. Phase contrast to observe iPSC morphology -tightly packed colonies with a high nucleus-to-cytoplasm ratio and pronounced nucleoli. Immunofluorescence image for pluripotency markers - OCT4 (red) and SSEA4 (green). Images taken with EVOS M7000 microscope using a 10x objective, scale bar = 275um.

Supplementary figure 2: Trilineage differentiation.

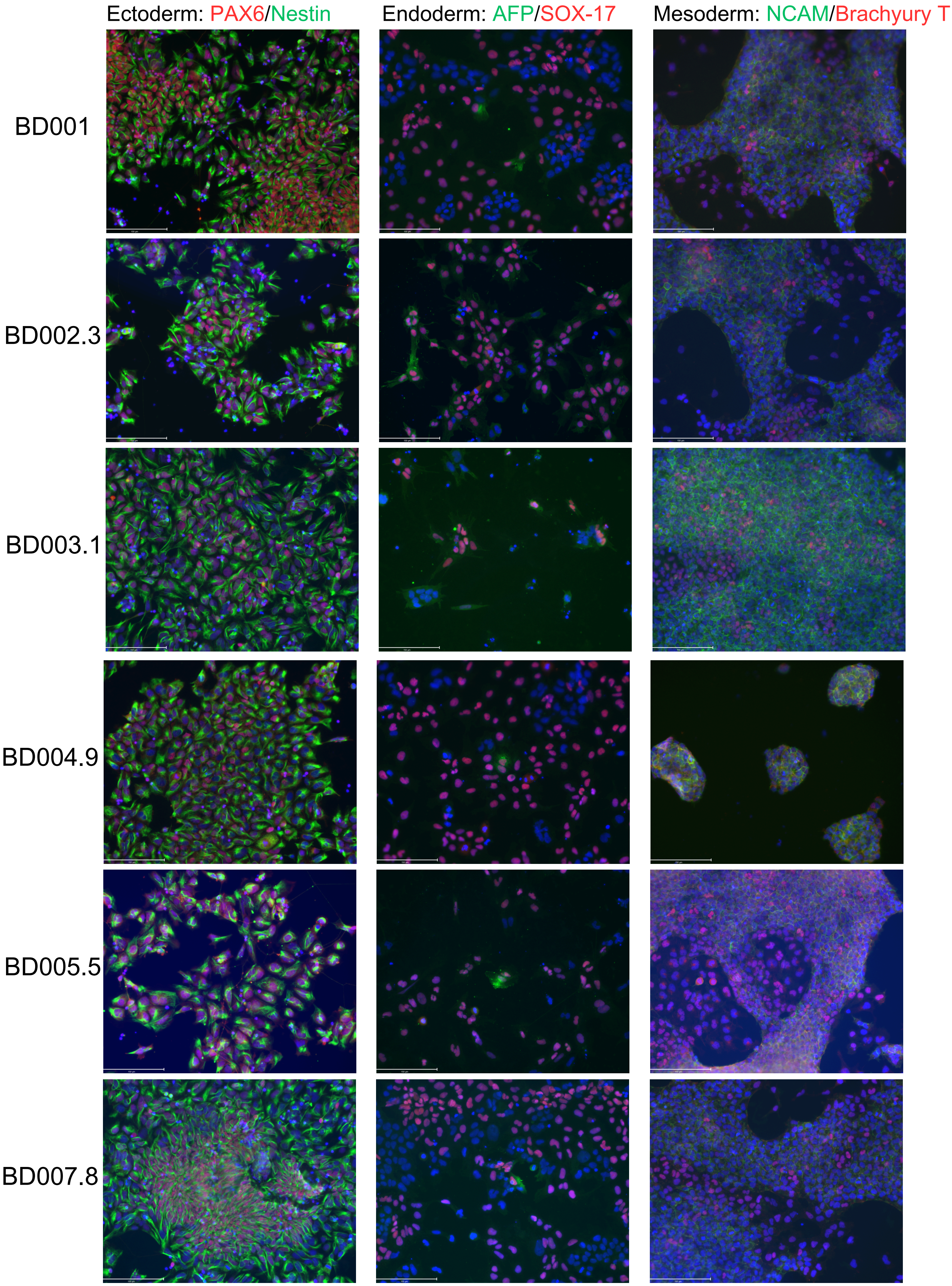

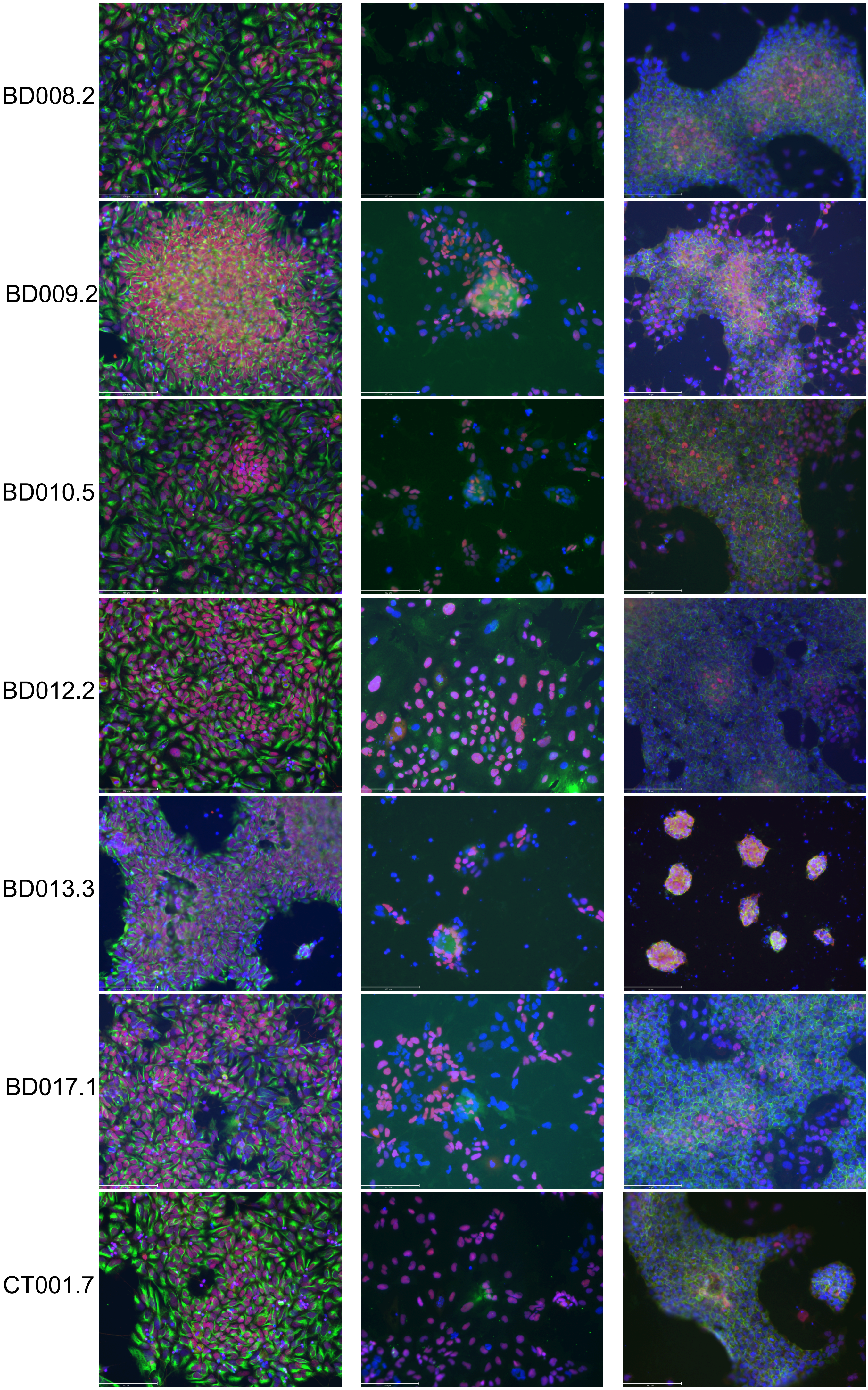
BD008.2
BD009.2
BD010.5
BD012.2
BD013.3
BD017.1
CT001.7

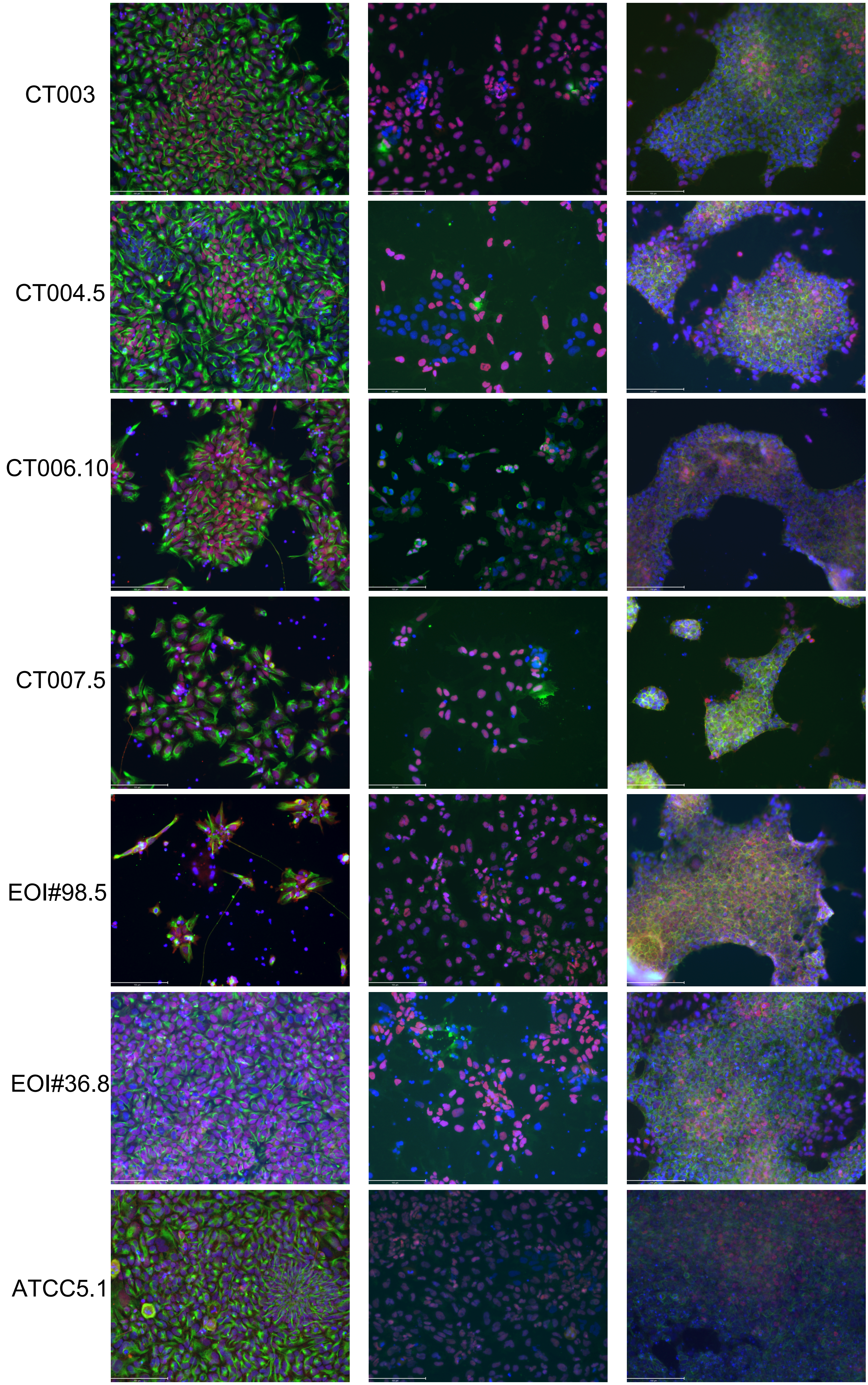
CT003
CT004.5
CT006.10
CT007.5
EOI#98.5
EOI#36.8
ATCC5.1

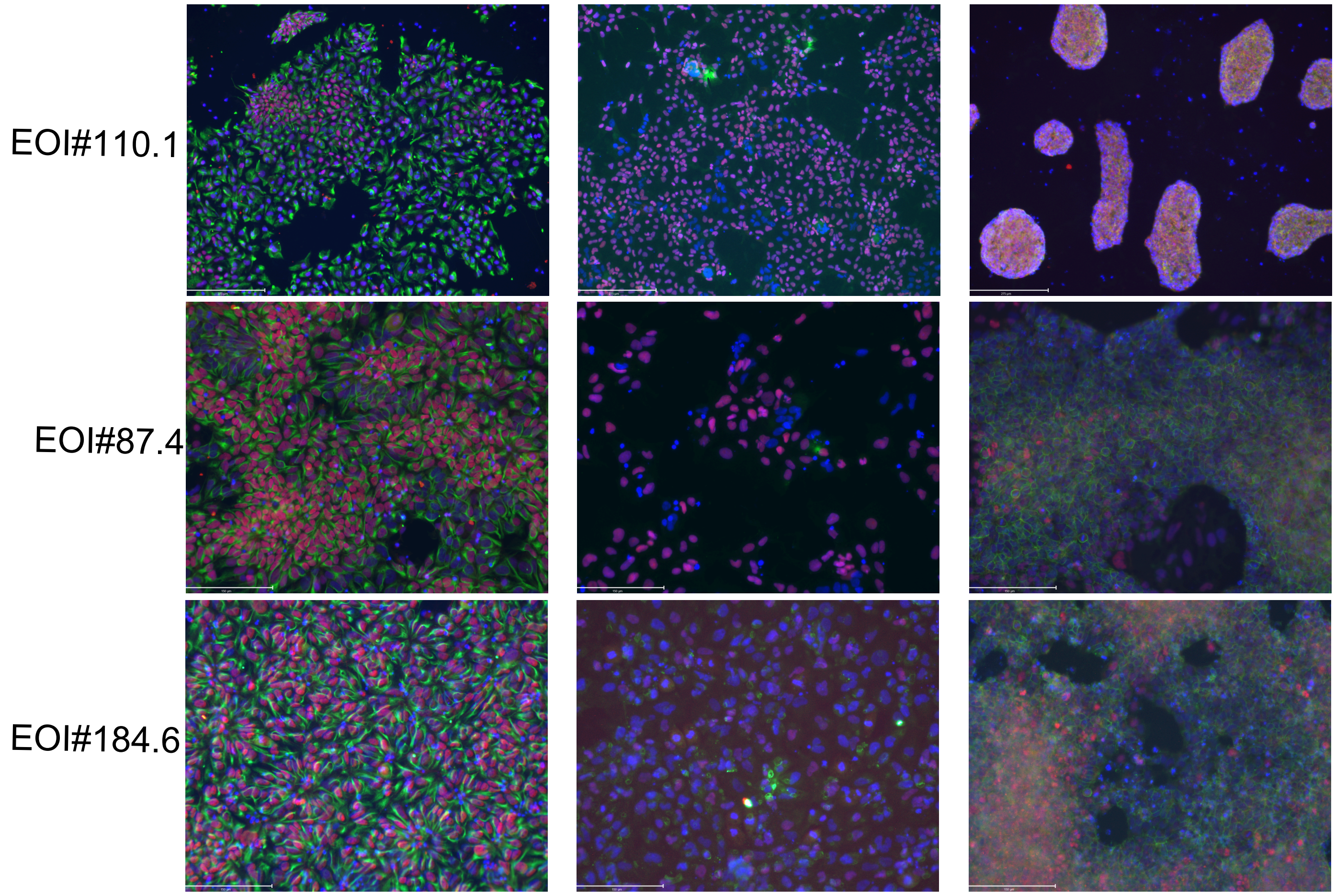


Images of trilineage differentiation for the 24 lines generated. Imunofloarescnece images for markers of the 3 germ layers: Ectoderm: PAX6 - red, Nestin - green; Endoderm: AFP - green and SOX17 - red; Mesoderm: NCAM - green and BrachuryT - red. Images were taken using the EVOS M7000 microscope using a 10x objective, scale bar = 150um.

Supplementary figure 3: Cortical networks

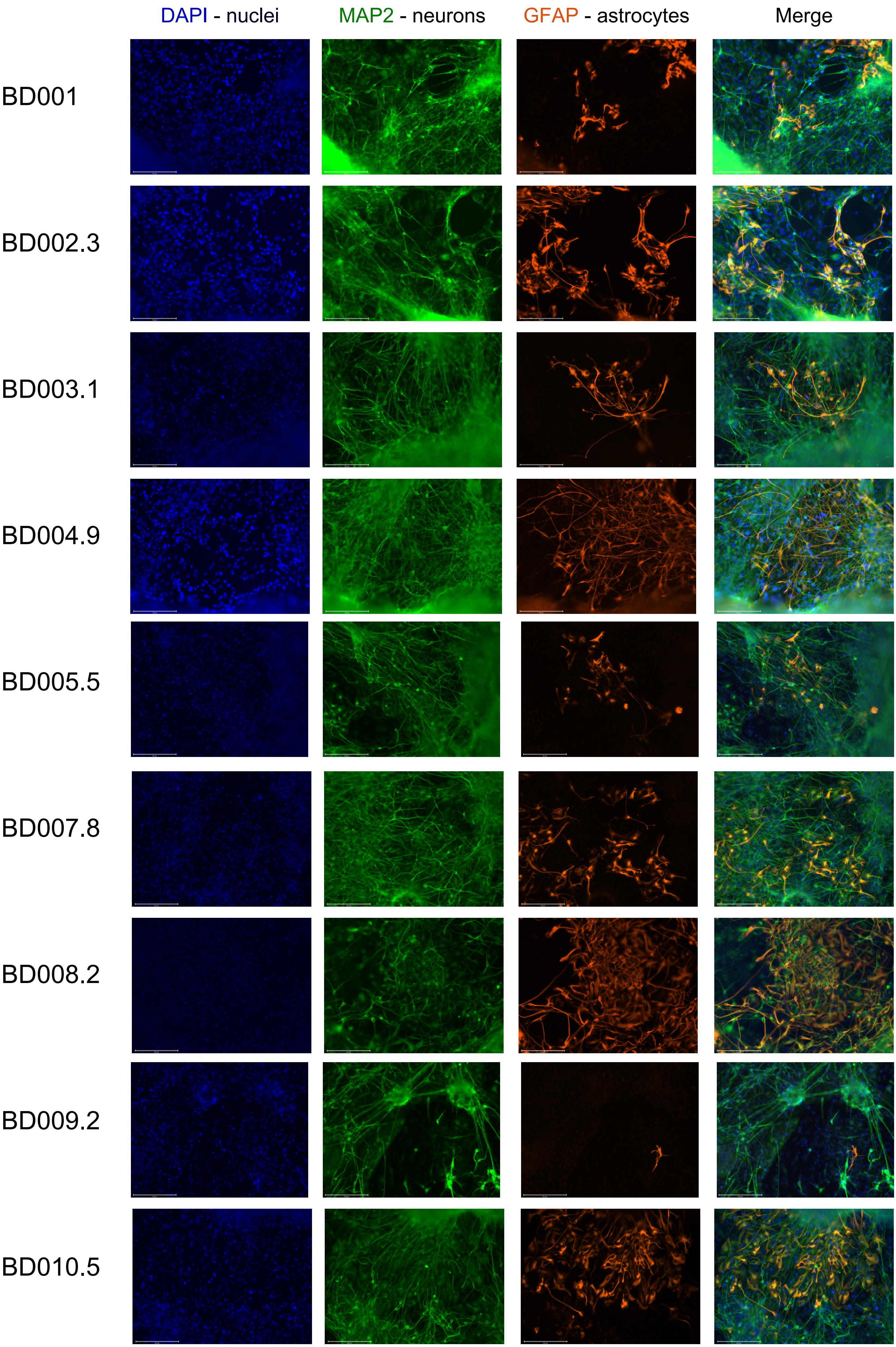

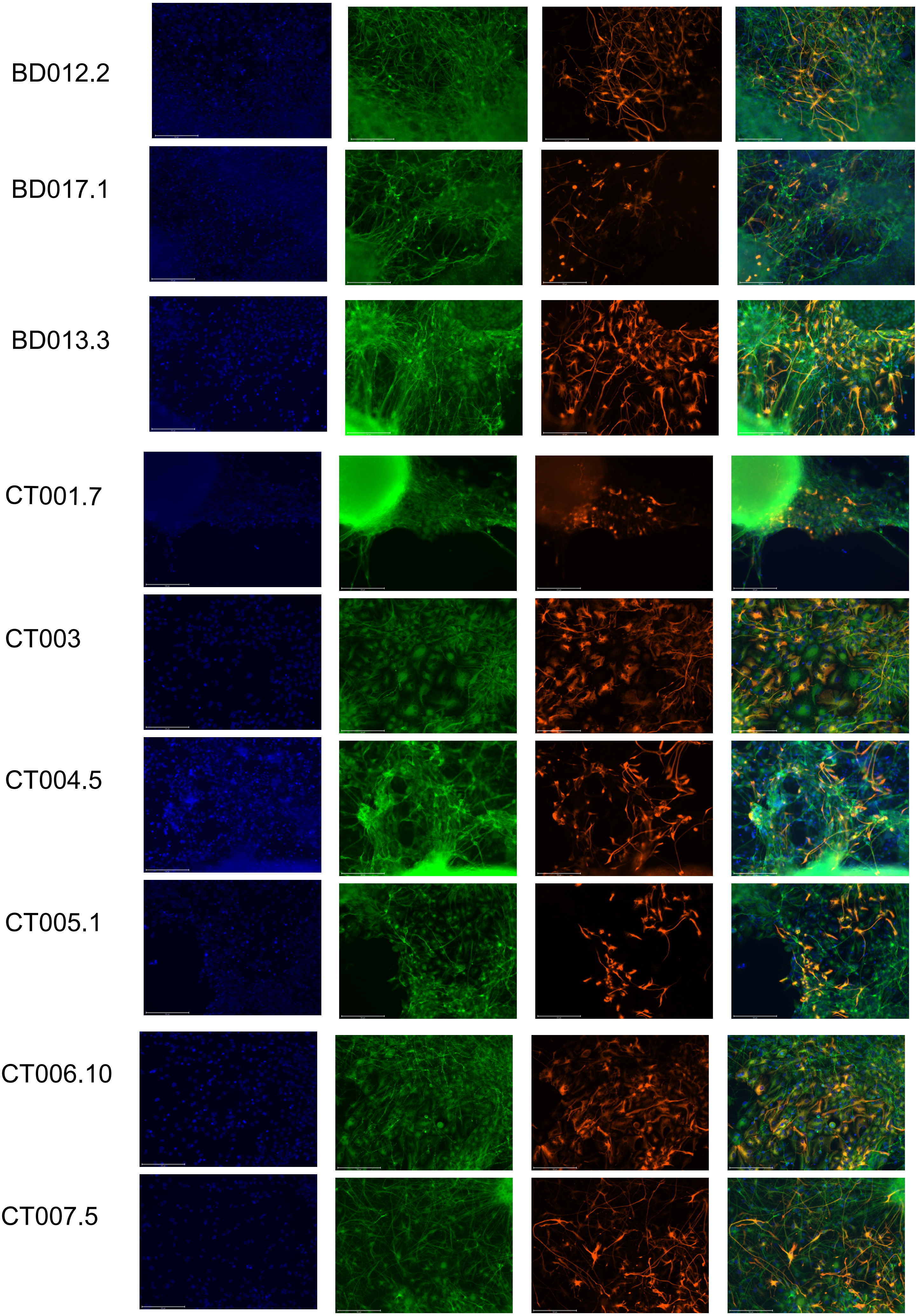

BD012.2
BD017.1
BD013.3
CT001.7
CT003
CT004.5
CT005.1
CT006.10
CT007.5

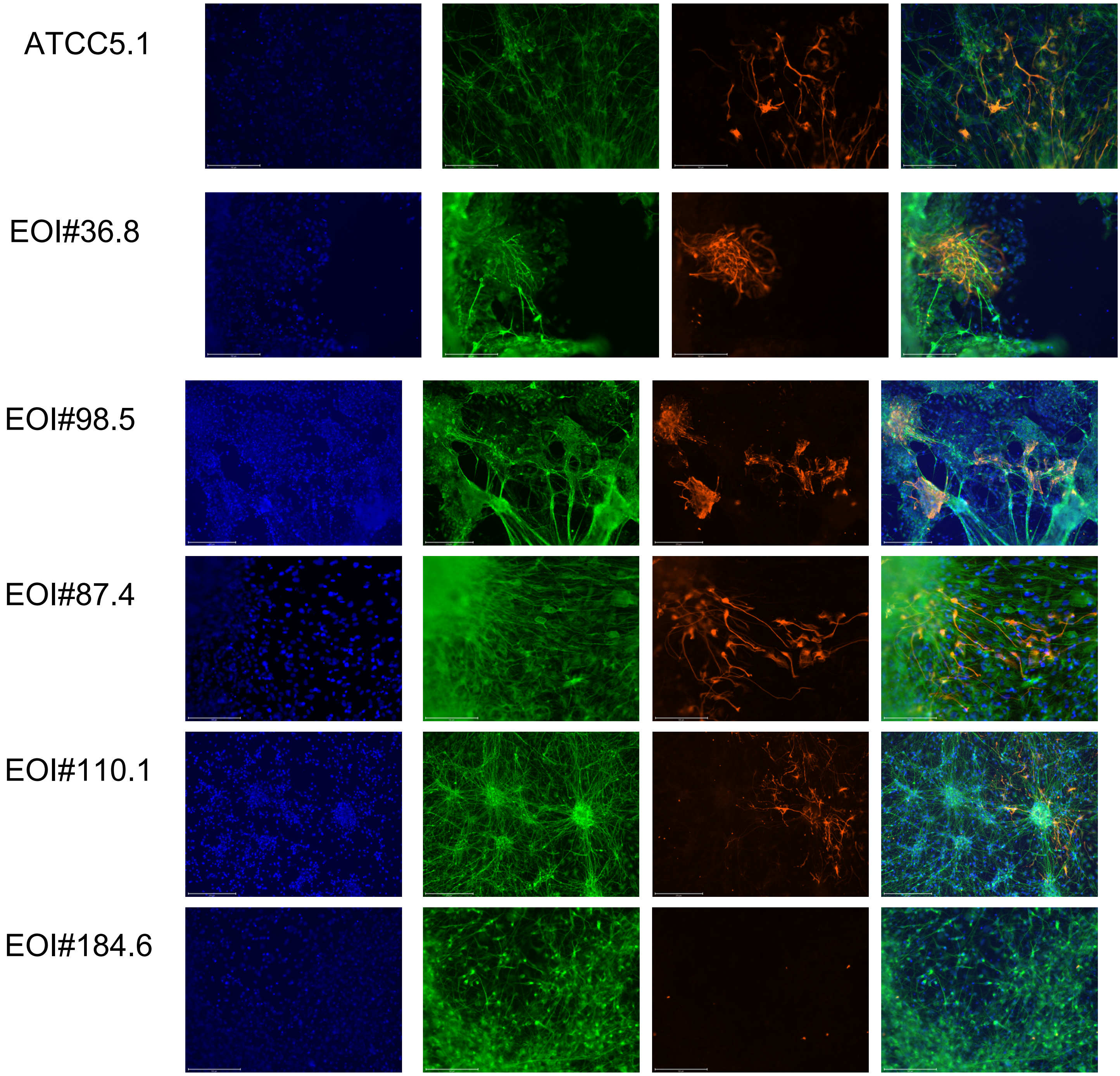


Immunofluorescence images of 24 lines differentiated into neurons (MAP2 - green) and astrocytes (GFAP - orange or purple). Images taken with EVOS M700 microscope with 20x objective, scale bar = 150um. Images are not processed, except to add a scale bar.
